# Stability and dynamics of the human gut microbiome and its association with systemic immune traits

**DOI:** 10.1101/2020.01.17.909853

**Authors:** Allyson L. Byrd, Menghan Liu, Kei E. Fujimura, Svetlana Lyalina, Deepti R. Nagarkar, Bruno Charbit, Jacob Bergstedt, Etienne Patin, Oliver J. Harrison, Lluís Quintana-Murci, Darragh Duffy, Matthew L. Albert, The Milieu Intérieur Consortium

**Author notes:** Correspondence (A.L.B.), (D.D.).

## Abstract

Analysis of 1,363 deeply sequenced gut microbiome samples from 946 healthy donors of the Milieu Intérieur cohort provides new opportunities to discover how the gut microbiome is associated with host factors and lifestyle parameters. Using a genome-based taxonomy to achieve higher resolution analysis, we found an enrichment of *Prevotella* species in males, and that bacterial profiles are dynamic across five decades of life (20-69), with Bacteroidota species consistently increased with age while Actinobacteriota species, including *Bifidobacterium*, decreased. Longitudinal sampling revealed short-term stability exceeds inter-individual differences; however, the degree of stability was variable between donors and influenced by their baseline community composition. We then integrated the microbiome results with systemic immunophenotypes to show that host/microbe associations discovered in animal models, such as T regulatory cells and short chain fatty acids, could be validated in human data. These results will enable personalized medicine approaches for microbial therapeutics and biomarkers.

## Introduction

Microbial therapeutics, including fecal microbiota transplants, bacterial consortia and probiotics are increasingly being tested in patients with *Clostridium difficile* infections and other gastrointestinal (GI) disorders (Allegretti et al., 2019), including inflammatory bowel disease, and more recently non-gastrointestinal indications such as autism (Kang et al., 2019) and cancer (Mullard, 2018). While the impact of microbial therapies on the recipients’ immune system has been characterized in animal models for several of these indications; in humans, the specific ways in which microbial therapies impact the recipient’s immune system are poorly understood. In parallel to microbial therapeutics, microbial signatures are being evaluated as a novel class of biomarkers, applied for stratification of efficacy and safety in clinical trials across multiple indications (Ananthakrishnan et al., 2017; Dubin et al., 2016). Notably, this rapid increase in microbial therapeutics and biomarkers justifies a careful reevaluation of the factors influencing an individual’s personal gut microbiome over time, as well as how gut microbes associate with systemic immunity in the steady state.

In this paper, we present a comprehensive assessment of the gut microbiome of 946 well-defined heathy French donors from the Milieu Intérieur (MI) consortium with 1,363 shotgun metagenomic samples. Designed to study the genetic and environmental factors underlying immunological variance between individuals, the MI consortium is comprised of 500 women and 500 men evenly stratified across 5 decades of life from 20 to 69 years of age, for whom extensive metadata, including demographic variables, serological measures, dietary information, as well as systemic immune profiles, are available (Patin et al., 2018; Thomas et al., 2015).

To build on the findings of several landmark microbiome studies (Falony et al., 2016; Human Microbiome Project, 2012; Zhernakova et al., 2016), many of which relied on an older reference library for taxonomic classification of microbial sequence reads (Truong et al., 2015), we leveraged an expanded set of reference genomes with a novel taxonomy that corrects many misclassifications in public databases to discover new biological insights particularly around age and sex (Parks et al., 2018). Notably, an independent dataset was used for replication of many of the findings (Zeevi et al., 2015). Longitudinal samplings from half the donors were used to expand on previous work that individuals are more similar to themselves over time, as compared to others (Costello et al., 2009; Flores et al., 2014; Mehta et al., 2018), and to identify factors that influence an individual’s stability over a two week time period. Using this improved annotation, we relate microbial composition to systemic immune profiles to uncover novel bacteria/immune interactions which can be experimentally tested and ultimately leveraged to design a microbial therapeutic with a desired immunogenicity. Overall this study utilizes the increased availability of reference genomes, granularity afforded by deep shotgun metagenomic sequencing, and statistical power of a large, well-characterized cohort to provide new insights into host/bacteria biology that will enable personalized medicine approaches for microbial therapeutics and biomarkers. In addition, the rich metadata and 1,000 plus deep shotgun metagenomic samples provided here will be a valuable resource upon which future microbiome studies can test and build new computational tools as well as generate and test new hypotheses.

## Results

### The Genome Tree Database improves taxonomic resolution of *k*-mer based approaches

Historically, microbial sequencing efforts focused predominantly on a small number of organisms, often causes of nosocomial infections [Figure S1A,B]. By contrast, reference databases, including NCBI Genbank, are increasingly populated with genomic information of commensal microbes (Browne et al., 2016; Forster et al., 2019; Poyet et al., 2019; Zou et al., 2019). As genome reference databases expand, historical, microbiological-based taxonomic assignments do not reflect population level relationships inferred from genome sequencing. This is particularly problematic for *k*-mer based analyses which utilize sequence similarity between closely related genomes to infer which taxa are present (Nasko et al., 2018).

To overcome these issues, in lieu of the traditional NCBI taxonomy, we generated a custom reference database of 23,505 RefSeq genomes with Genome Tree Database (GTDB) taxonomies (See methods)[Table S1]. Briefly, GTDB is a bacterial taxonomy based on a concatenated protein phylogeny in which polyphyletic groups were removed and taxonomic ranks were normalized on the basis of relative evolutionary divergence (Parks et al., 2018). The impact of this procedure was particularly prominent for species of the genus *Clostridium*, which were split into 121 unique genera spanning 29 families (Parks et al., 2018). This could be especially meaningful for analysis of gut microbiome samples as *Clostridium* species are a prevalent community members and often emerge in association studies.

The Refseq sequences and taxonomic tree from the GTDB were used to build a reference database for the *k*-mer based program Kraken2 (Wood and Salzberg, 2014) and read-reassignment step Bracken (Lu et al., 2017). This custom Kraken2/GTDB pipeline was applied to 1,363 quality-controlled samples from 946 MI donors [Figure S1C-F, Table S2,3], and compared using both the marker gene based tool Metaphlan2 (Truong et al., 2015) and Kraken2 with the same 23,505 reference genomes using their original NCBI taxonomies [Figure S2]. Consistently, more bacterial taxa were identified per sample with Kraken2 than Metaphlan2, a result of the updated reference database and higher sensitivity of this *k*-mer based approach (McIntyre et al., 2017) [Figure S2A-C]. Between the two Kraken databases (GTDB and NCBI), richness varied depending on how taxa were re-distributed by GTDB. For example, GTDB split 2,397 NCBI genera into 3,205, while it collapsed 18,795 NCBI species into 13,446 [Figure S2A,D]. Despite finer level differences, the overall distribution of phyla across the 3 approaches was similar [Figure S2E], indicating that Kraken2/GTDB pipeline results would be consistent with previous analyses. As such, a combination of *k*-mer based read assignment and genome-based taxonomy allows higher resolution analysis of shotgun metagenomic samples.

### Variable gut microbiomes in a restricted geographical region

To complement our optimized taxa-based approach and further utilize the resolution afforded by shotgun metagenomic sequencing, we applied HUMANn2 to identify the functional potential of microbial pathways present in the MI samples (Franzosa et al., 2018) [Table S4]. Utilizing both the Kraken2/GTDB and HUMANn2 pipelines, we identified a broad range of diversity across the 946 individuals in this geographically-restricted cohort of healthy French adults. This diversity was observed in terms of metabolic pathway richness (282 ± 40), species richness (353 ± 45), as well as Shannon diversity (3.9 ± 0.36), which accounts for both richness and evenness [Figure 1A-C, Table S2]. Across donors, our GTDB pipeline confirmed Firmicutes and Bacteroidota (formerly Bacteroidetes) as the most abundant phyla in the gut, but enabled distinction among the original Firmicutes phyla, which was further divided into 12 distinct categories; Firmicutes, Firmicutes_A, Firmicutes_B, … Firmicutes_K [Table S1]. Notably, throughout the GTDB, the group containing type material (if known) kept the original unsuffixed name. Of those, 7 were present in this cohort with Firmicutes_A the most abundant, followed by Firmicutes and Firmicutes_C [Figure 1D, Table S3], highlighting the finer granularity, even at the phylum level, provided by GTDB-based taxonomic calls. Subsequent application of the Bray-Curtis distance metric, as a means to assess species presence/absence in addition to relative abundance across donors, demonstrated that samples fell along a gradient defined by the relative abundances of Firmicutes_A and Bacteroidota [1st dimension of MDS projection, Figure 1E].

**Figure 1.**
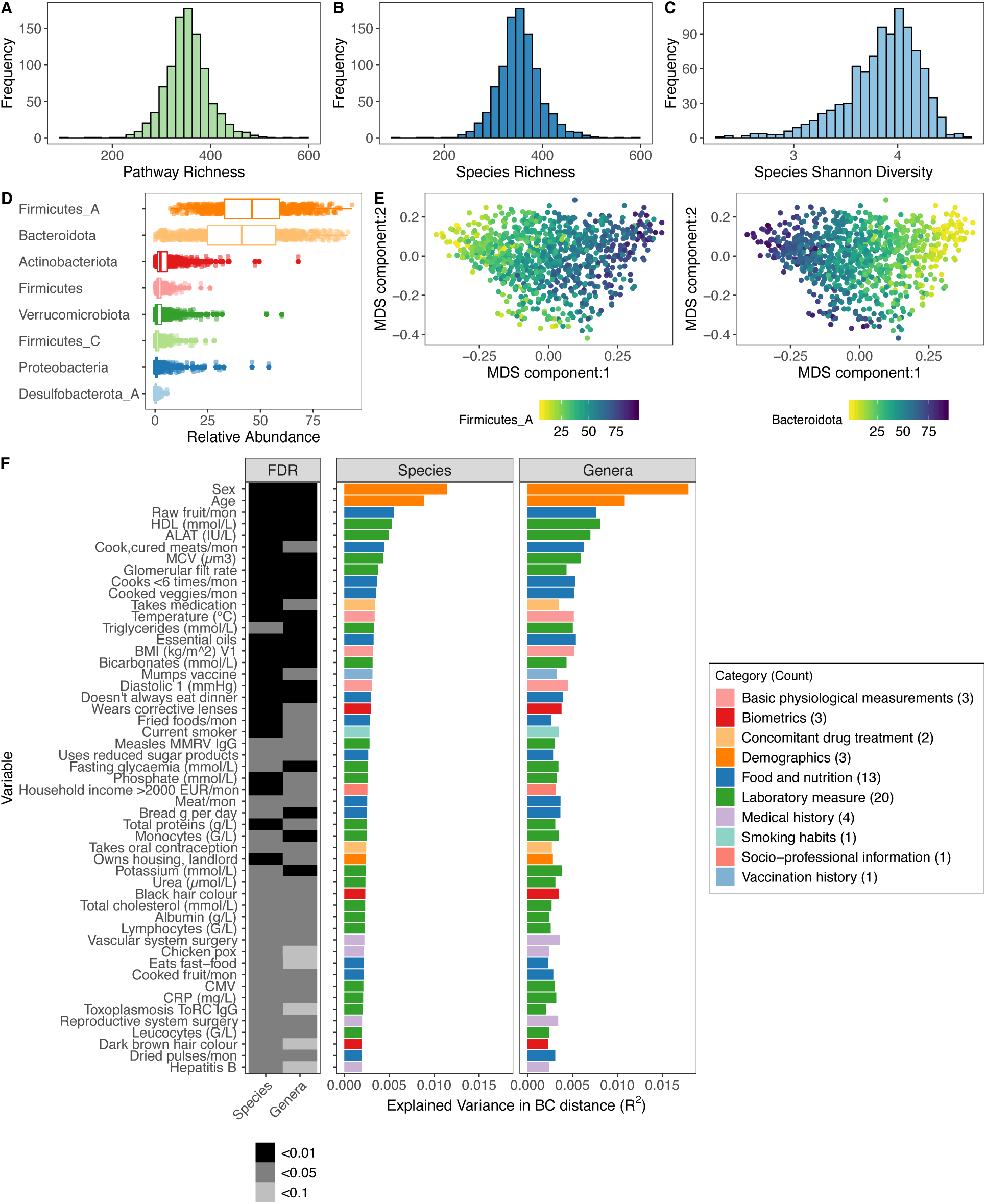
Inter-individual variation of bacterial composition is associated with many factors. (A) Distribution of pathway richness across donors. (B) Distribution of species richness across donors. (C) Distribution of species Shannon diversity index across donors. (D) Boxplots of the top 8 phyla. Each dot corresponds to 1 donor. Firmicutes_A, Firmicutes_C, and Firmicutes were split into unique phyla by the Genome Tree Database (GTDB). (E) Multidimensional scaling (MDS) plots of Bray-Curtis distance of bacterial species composition. Ordination was driven by the top two phyla Firmcutes_A and Bacteroidota. Each dot corresponds to 1 donor while color indicates relative abundance of each phyla. (F) In total, 51 factors (Benjamini & Hochberg FDR < 0.05) were associated with inter-individual variation of the gut microbiome. The bar plots indicate the amount of interindividual variance explained by each factor for the species and genera level Bray-Curtis (BC) distance. Variables are ordered by the % variance explained in Kraken species. Colors of the bars correspond to the broad metadata category. The rectangles to the left indicate the statistical strength, as measured by FDR, of the association. For each variable, samples with NA values were excluded. For all these analyses, 1 sample per donor was used, when available Visit 1, if not Visit2. See also Figures S1, S2, S3, S4, and S5 and Tables S1, S2, S3, S4, S5, S6, and S7.

Using this extensively characterized cohort we explored how 154 metadata variables [Figure S3, Table S5,6], including 42 laboratory measurements and 43 dietary variables, contributed to overall bacterial community composition. We identified 51 variables (33% of total) that associated with Kraken2-GTDB species (multivariate PERMANOVA test FDR <0.05); all of which replicated at the genera-level (multivariate PERMANOVA test FDR <0.1) [Figure 1F, Table S7]. The top contributors were age and sex with lesser contributions from diet such as consumption of raw fruit, cooked, cured meats, as well as frequency of fast food consumption, in line with previous reports of 16S rRNA analyses from this cohort (Partula et al., 2019; Scepanovic et al., 2019). Notably, sex and age were associated with 24 and 44 of the other metadata variables respectively, which confounds our ability to dissociate the individual effects of these variables on microbial community composition [Figure S4]. In total, these factors explained less than 10% of population variability, indicating that the majority of variance in community composition remains unexplained. Drawbacks of this analysis are the absence of Bristol stool score, a measure of stool consistency, and levels of chromogranin A, a protein secreted by enteroendocrine cells, the factors most associated with community composition in previous European cohorts (Falony et al., 2016; Zhernakova et al., 2016). Although genetic data were also available for these donors, they were not considered here based on previous analyses that the effects of host genetics on microbiome are minimal in this (Scepanovic et al., 2019) and other cohorts (Rothschild et al., 2018) in part due to the small population sizes by GWAS standards (Goodrich et al., 2017).

Notably, in this healthy cohort medication usage was low, with only 28% of individuals (n=266) taking medication of any kind. Of all medications, only oral contraception was taken by more than 10% of participants (n=111). In pre-menopausal women, oral contraception was taken by 36% (110/303) and explained 0.005% of the variance (p-value = 0.002). In contrast, relatively common medications, beta-blockers and proton pump inhibitors were taken by only 7 and 4 individuals respectively. Despite this, medication usage was a significant albeit minor contributor (R^2^ = 0.003) to microbial community composition, highlighting how xenobiotics can, and do influence the gut microbiome (Jackson et al., 2018; Maier et al., 2018).

### MI bacterial profiles are comparable to those from Israeli donors

In order to determine whether MI bacterial profiles were unique to this population, or comparable with other non-European healthy cohorts, we ran our Kraken2/GTDB pipeline on 1,161 samples from 851 Israeli donors originally published by Zeevi et al (Zeevi et al., 2015) [Table S8], for whom age, sex, and body mass index (BMI) were provided [Figure S5A,B]. After accounting for sequencing depth (mean read count MI: 14.2 million vs. Zeevi 13.5 million), we found that richness across taxonomic levels was consistently elevated in the MI samples, even though the percentage of unmapped reads was comparable (MI 42.2% versus Zeevi 39.5%) [Figure S5C-E]. More specifically, we identified on average 75 more species in samples from the MI donors compared to those from the Zeevi cohort. In addition to potential technical and lifestyle reasons, this discrepancy could reflect the stricter inclusion and exclusion criteria, and thus the greater overall health of the MI donors (Thomas et al., 2015; Zeevi et al., 2015).

On the whole, community composition including relative taxa abundances and beta diversity was consistent across both cohorts [Figure S5F-H]. Notably however, the contributions of age and sex to community composition were almost 2 times greater in MI than Zeevi (Age: R^2^ 0.0088 vs. 0.0038, Sex: 0.011 vs. 0.0068)[Figure S5I], highlighting how stratification of age and sex in the MI cohort provided enhanced statistical power to identify new associations [Figure S5A,B]. Despite the geographic and cultural distinctions between these cohorts, our findings demonstrate a comparable makeup of the gut microbiome. This allowed us to utilize the Zeevi samples as a replication cohort to demonstrate the reproducibility of our findings in MI.

### *Prevotella* species are more abundant in male donors

Given that sex and age were the variables most strongly associated with bacterial community composition in healthy individuals, we leveraged the statistical power of the MI cohort to explore which taxa were differentially abundant between sexes and across decades of life. To identify bacteria differentially abundant between the 473 females and 473 males, we conducted DESeq2 analysis (Love et al., 2014) on 547 abundant species (prevalence > 5% and a mean relative abundance > 0.01%) [Table S9]. Of the 130 differentially abundant species (FDR < 0.05) [Figure 2A], 12 were more abundant in females, while 26 were more abundant in males with log2 fold change > 1 [Figure 2B,C]. In total, 17 out of 45 prevalent *Prevotella* species were more abundant in males compared to females, corresponding to a greater overall richness of *Prevotella* species in males [Figure 2D]. Similarly, in the Zeevi cohort, 25 species of *Prevotella* were more abundant in males [Figure S6, Table S10]. Notably, even when a species was significantly differentially abundant between sexes in only one cohort, the direction of this trend was also consistent in the other, indicating that higher *Prevotella* abundance in males compared to females is a biological phenomenon consistent across multiple species and populations. This information expands the findings of two previous studies, one that identified *Bacteroides*-*Prevotella* as broadly more abundant in males than females based on 16S rRNA-targeted oligonucleotide probes (Mueller et al., 2006) and another that found males were three times more likely to have an enterotype consisting of fewer *Bacteroides* and higher *Prevotella* (Ding and Schloss, 2014). Although the factors driving preferential colonization of *Prevotella* in males are unknown, from these data we could generate hypotheses surrounding the roles of gonadal hormones and microbial community composition.

**Figure 2.**
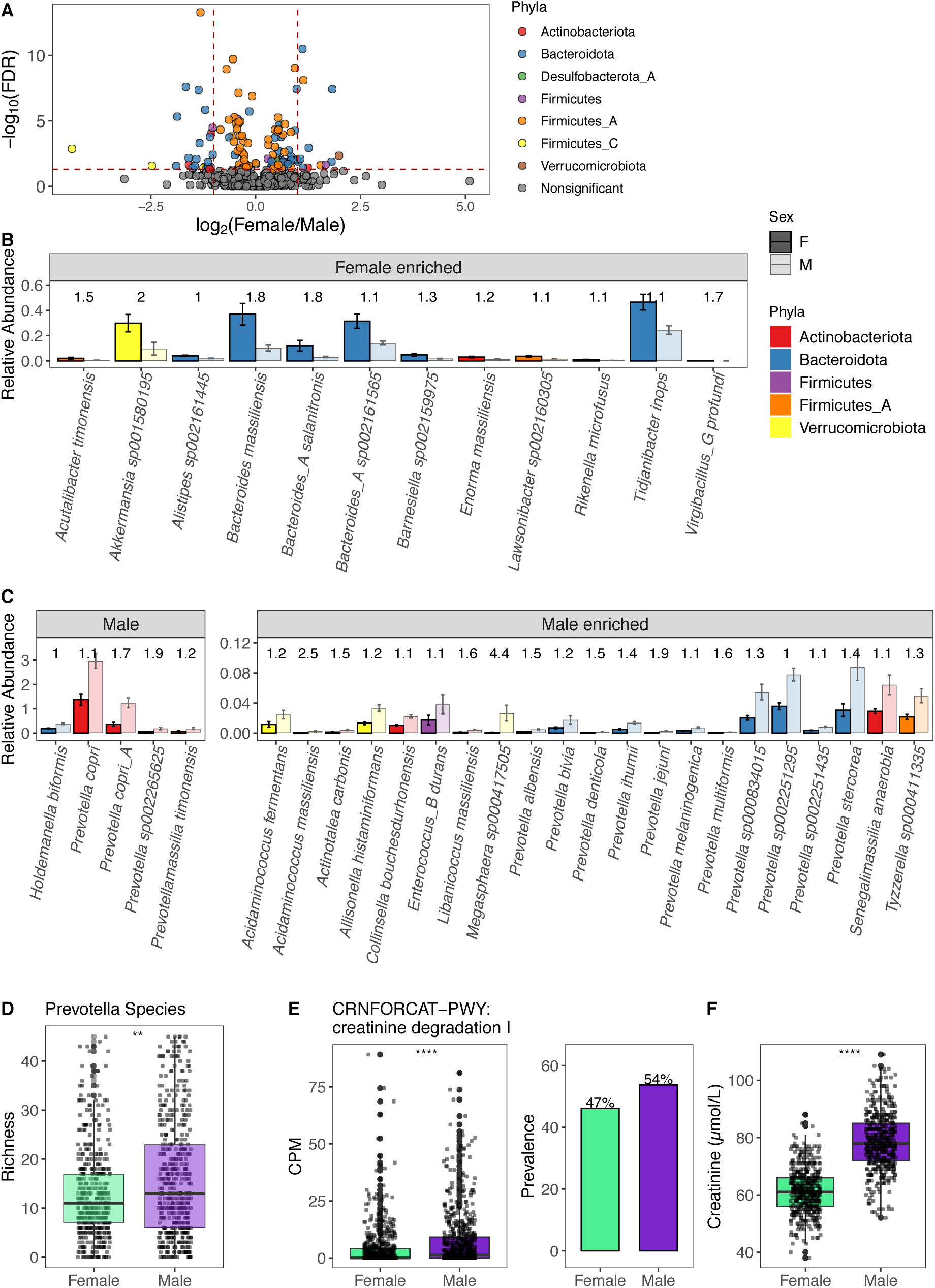
130 bacteria were differentially present between males and females. (A) Volcano plot of 547 abundant species, of which 130 were differentially abundant between males and females based on DESeq2 (FDR < 0.05). Each species is colored by its taxonomic phyla. (B, C) Bars indicate the mean relative abundance of the species in either sex. The left bars with a black border are for females, while those shaded with a gray border are for males. Error bars indicate the mean ± standard error. Absolute log2 fold change for each species is indicated above the bars. (B) 12 bacteria enriched in females with FDR < 0.05 and log2 fold change > 1. (C) 26 bacteria enriched in males with FDR < 0.05 and absolute log2 fold change > 1. (D) Boxplots show richness of *Prevotella* species in males and females. (E) Abundance measure in Counts Per Million (CPM) and prevalence of the MetaCyc pathway CRNFORCAT-PWY: creatinine degradation I in males and females. (F) Blood Creatinine levels in males and females. (D-F) ** p < 0.01, **** p < 0.0001 by Wilcoxon-Rank Sum. See also Figure S6 and Tables S9 and S11.

When considering metabolic pathways, we identified 54 (FDR < 0.05) differentially abundant between the sexes [Table S11]. Of those, the pathway CRNFORCAT-PWY: creatinine degradation I was the most strongly enriched in men [Figure 2E]. Biologically, this is consistent with men having greater blood levels of creatinine [Figure 2F]. Across both sexes but not in each individually, circulating creatinine levels were significantly associated with the abundance of the CRNFORCAT-PWY pathway (Both sexes Spearman rho = 0.087, p-value = 0.0014; men rho = 0.028, p-value = 0.47; women rho = -0.047, p-value = 0.22). Overall, this exemplifies how adaptation to utilize available nutrients may influence microbiome composition.

### The gut microbiome is dynamic across decades of life

The composition of the gut microbiome differs dramatically between newborns and adults, with the neonatal microbiome transitioning to a more adult-like state upon consumption of solid food and cessation of breast-feeding (Backhed et al., 2015; Stewart et al., 2018). Similarly, comparison of young adults, elderly individuals, and centenarians has identified differences in microbial community by culturing or 16S rRNA (reviewed by (An et al., 2018)). The design of the MI cohort provides a unique opportunity to explore how in the absence of underlying disease the gut microbiome is dynamic across the adult decades (20 to 69 years old).

In total, we identified 227 out of 547 abundant species that were differently abundant by donor age (Spearman correlations of age by taxa relative abundance, FDR < 0.05) [Table S12]. Notably, 3 of the top 5 phyla (Bacteroidota, Actinobacteriota, and Proteobacteria), experienced shifts in relative abundance across the decades [Figure 3A]. Transitions were most pronounced around 40-50 years old, a time span when many persons experience the preclinical stages of chronic diseases, and women begin to experience hormonal changes associated with onset of menopause; average age in this cohort 50 ± 4.2 years. Across phyla, the associations with age were conserved across sexes and cohorts [Figure S7A,B]. For example, relative abundances of Proteobacteria and Bacteroidota were primarily increased with age, while Actinobacteriota, including 17 species of *Bifidobacterium* were decreased with age [Figure 3B]. This gradual decline of *Bifidobacterium* was true in terms of both relative abundance as well as presence/absence [Figure 3C,D]. Notably, *Collinsella* species that were positively correlated with *Bifidobacterium* were also decreased with age [Figure 2B, Figure S7C].

**Figure 3.**
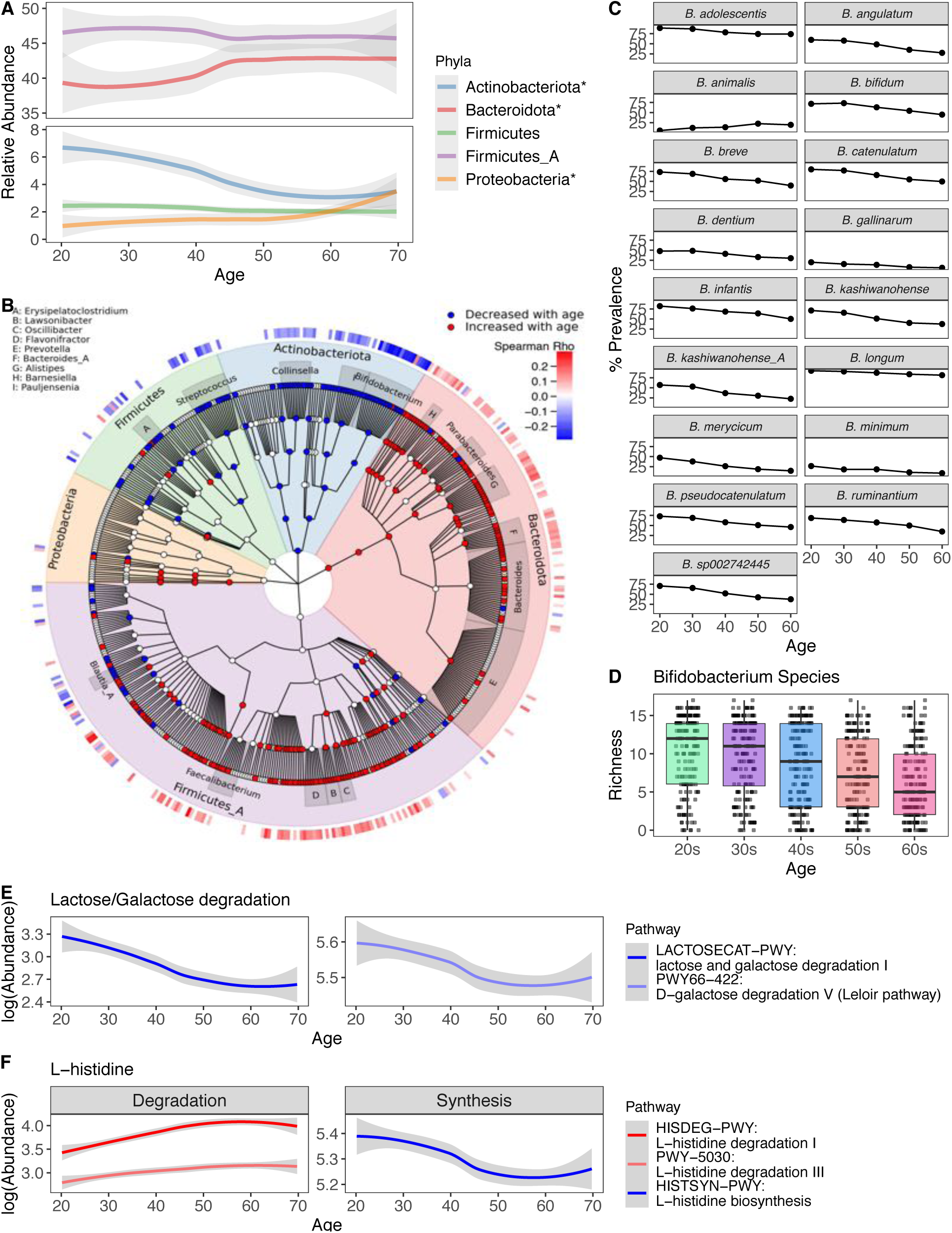
Bacterial profiles are dynamic across decades of life. (A) Abundance of the top 5 most abundant phyla across decades of life. Curves show 95% confidence intervals and were modeled with LOESS regression. Stars in the figure legend indicate those phyla statistically associated with age. (B) GraPhlAn taxonomic tree of the 508 species in the top 5 phyla found at significant prevalence and abundance across all donors. Relative abundance of taxa in red were decreased with age, while those in blue increased. Association between taxa relative abundance and age was determined by Spearman correlation (FDR < 0.05). The heatmap on the outer ring indicates the strength of the correlation (Spearman Rho). Genera with at least 5 species associated with age are labeled. (C) Lines indicate the percent prevalence of 17 *Bifidobacterium* species across the different decades. Only *Bifidobacterium* species significantly associated with age are shown. (D) Boxplots show richness of *Bifidobacterium* species in donors grouped by their age. (E) Abundance (CPM) of lactose/galactose degradation pathways decreased with age. Curves show 95% confidence intervals and were modeled with LOESS regression. (F) Abundance (CPM) of L-histidine degradation/synthesis pathways associated with age. Curves show 95% confidence intervals and were modeled with LOESS regression. See also Figure S7 and Tables S12 and S14.

The decline in *Bifidobacterium* prominence with age is particularly interesting in light of *Bifidobacterium* being the dominant bacteria in newborns, gradually decreasing as infants cease breastfeeding (Stewart et al., 2018). The association of *Bifidobacterium* and old age indicates that the loss of *Bifidobacterium* occurs not only in infants, but continues throughout adulthood (An et al., 2018; Biagi et al., 2016; Biagi et al., 2010; Kato et al., 2017; Mueller et al., 2006). Similar to our findings associating *Prevotella* and sex [Figure S6B], we built on previous findings and revealed the trend was consistent across species within the genera and across cohorts [Figure S7D, Table S13], highlighting how the phenomenon is intrinsic to this species. Notably, the only exception, *Bifidobacterium animalis,* is a common probiotic-associated strain (Turroni et al., 2009), rather than a persistent colonizer. Moving forward, comparative genomic analyses between these different species could reveal features associated with colonization in older adults.

We then focused our attention on 364 prevalent microbial pathways (prevalence > 5%) and identified 108 that correlated with age [Table S14], of which 31 were increased and 77 were decreased [FDR < 0.05], including several lactose and galactose degradation pathways [Figure 3E]. Lower levels of lactose/galactose degradation may explain increased lactose intolerance in older adults and presents a possible opportunity for microbial therapeutic intervention (Gingold-Belfer et al., 2019; Savaiano et al., 2013). Notably in this cohort, the abundance of these pathways was not associated with consumption of dairy products, e.g. milk, cheese, and yogurt (Spearman p-values >0.3).

Other pathways associated with age were related to L-histidine. In this case, pathways for L-histidine biosynthesis were decreased with age while those for degradation were increased [Figure 3F], indicating that gut L-histidine levels may be decreased in older adults which could lead to an altered immune state as L-histidine metabolites have been demonstrated to influence colonic inflammation (Gao et al., 2017). Overall, understanding the multitude of microbial associations with age will become increasingly important as microbial therapeutics are being evaluated as treatments for individuals of all ages.

### Short term stability is variable across donors

To complement our cross-sectional study of the microbiome across the decades, we leveraged longitudinal sampling of roughly half the cohort (n = 413) to study short-term (17 ± 3.3 days) dynamics within an individual in the absence of antibiotic exposure. By comparing species Bray-Curtis distances within and between individuals, we found that in the short-term intra-individual differences were less than the inter-individual ones [Figure 4A]. This is consistent with previously published findings (Costello et al., 2009; Flores et al., 2014; Mehta et al., 2018), and also reflected the analysis of relative abundance and metabolic pathways presence/absence [Figure 4A]. Notably, differences between donors were less dramatic at the pathway level, reflecting the more conserved nature of annotated metabolic pathways versus species profiles across individuals (Human Microbiome Project, 2012).

**Figure 4.**
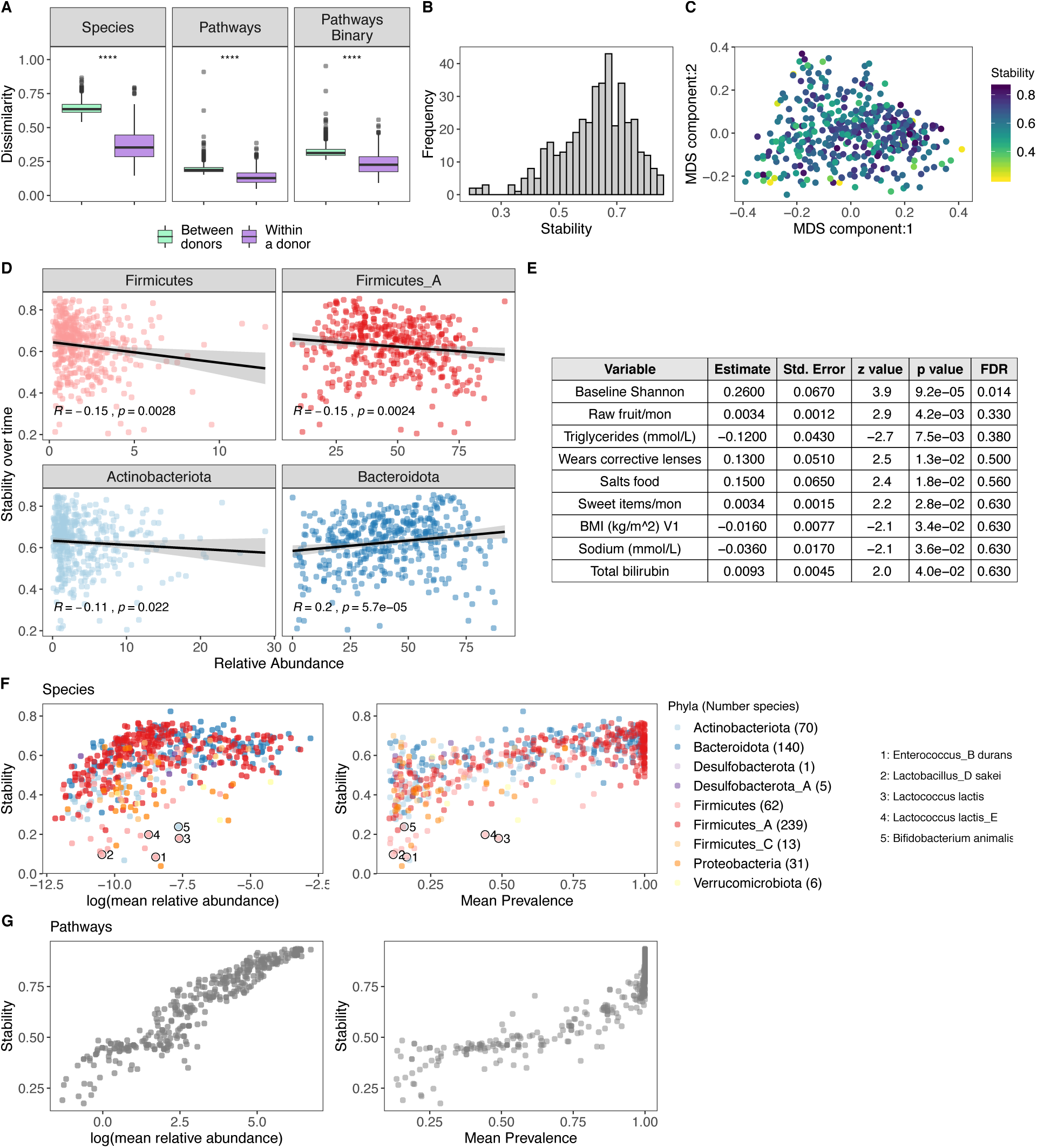
Bacterial profiles are stable over the short-term, however the degree of stability is variable across donors. (A) Boxplots of species Bray-Curtis distances (left), pathway Bray-Curtis distances (middle), and pathway binary Jaccard distances (right) between donors and within a donor over time (n= 413, time between samples= 17 ± 3.3 days), 1 = samples are completely different, 0 = samples are identical. **** p-value < 2.2e-16 by Wilcoxon-Rank Sum. (B) Histogram of 413 donors’ species stability (1-Bray-Curtis). (C) Multidimensional scaling (MDS) plot of Bray-Curtis distance of bacterial species composition. Each dot corresponds to 1 donor who had a second sample. Color indicates longitudinal stability of that donor (1 - within sample Bray-Curtis). (D) Scatter plots show the top phyla associated with stability. Each point corresponds to 1 donor with a longitudinal sample (n= 413). Colors correspond to those in F. Trend lines show 95% confidence intervals and were modeled with lm. Stats based on Spearman correlation. (E) Results of the series of GLMM fits aimed at identifying factors associated with intra-individual stability. Only factors with p.value < 0.05 are shown. (F) Scatter plots show the stability of individual species (1 - Bray-Curtis) by their mean baseline relative abundance and prevalence. Each point corresponds to a bacterial species and is colored by the species Phyla. (G) Scatter plots show the stability of individual pathways (1 - Bray-Curtis) by their mean baseline relative abundance and prevalence. Each point corresponds to a pathway. See also Tables S15, S16, S17, and S18.

Although stability was the norm, the degree of stability (quantified as 1 - Bray Curtis distance (Mehta et al., 2018)) was variable across the 413 donors [Figure 4B,C], and as such we investigated which microbial and metadata features underlie this personalized stability trait. Using Spearman correlations, we identified the phyla Firmicutes, Firmicutes_A and Actinobacteriota as enriched at baseline in the less stable donors, while Bacteroidota was higher in donors with greater stability over time (Flores et al., 2014) [Figure 4D, TableS15]. This is consistent with observations that spore forming bacteria, including many Firmicutes species, are intrinsically less stable (Kearney et al., 2018). Using generalized linear models, we found BMI and Triglycerides levels were negatively associated with stability, while conversely, consumption of sweet items (e.g. chocolate, sweets, honey, jam) and raw fruit were positively associated [Figure 4E, Table S16], concordant with diet being a key determinant of human gut microbiome variation (Johnson et al., 2019). While the previous results were only marginally significant (p-value < 0.05), consistent with previous findings (Flores et al., 2014; Mehta et al., 2018) and ecological theory (McCann, 2000; Schindler et al., 2010; Tilman, 1999), baseline species Shannon diversity was positively associated with stability (FDR = 0.014) [Figure 4E], i.e. individuals with more diverse communities were more resilient to change than individuals with lower diversity.

We then applied the Bray-Curtis metric to calculate stability (1 – BC) of individual species and pathways [Table S17,18], as previously done (Faith et al., 2013; Franzosa et al., 2015). For both species and pathways, we found that their stability was strongly associated with their mean abundance and prevalence across donors [Figure 4F, G]. For example, the Firmicutes *Enterococcus_B_durans* had a mean abundance of 0.021%, prevalence 7%, and the third lowest stability 0.085. Additionally, many species known to be present in yogurt and probiotics were also highly unstable, e.g. *Lactobacillus_D sakei*, *Lactococcus lactis*, and *Bifidobacterium animalis* [Figure 4F] (Fijan, 2014); in agreement with previous observations that probiotics often face colonization resistance (Zmora et al., 2018). Moving forward, these data can be leveraged to prioritize microbial pathways/species that will make reliable biomarkers as well as persistent colonizers if incorporated into a microbial therapeutic.

### Bacterial species and microbial pathways are associated with systemic immunophenotypes

The microbiota has been shown to play a major role in the development, education and function of the mammalian immune system, largely through the use of mouse models (Belkaid and Harrison, 2017). However, there remains a gap in our knowledge concerning the association of microbial signatures and immune activation in healthy humans. Having established high resolution bacterial profiles, we leveraged immunophenotyping of matched peripheral blood samples to explore how gut bacterial species/pathways were associated with 86 systemic immune cell profiles (Patin et al., 2018). Of these immunophenotypes, 50 related to adaptive immunity and 36 with innate immune cell subsets [Figures S8,9, Tables S19,20]. After correcting for age, sex, cytomegalovirus (CMV) status, smoking, genetics, and batch, which were previously shown to associate with immune profiles (Patin et al., 2018), we applied multivariate linear models to associate 547 bacterial species (prevalence >5%, average abundance > 0.01%) and 350 metabolic pathways (prevalence > 20%) with the 86 immunophenotypes.

To evaluate the applicability of this approach, we initially investigated microbial species associated with circulating regulatory T (T_reg_) and TCRγδ^+^ T cell populations. Experimentally in animal models, unique microbial species have been previously demonstrated to engage these distinct cellular subsets (Belkaid and Harrison, 2017). In total, 37 pathways (FDR ≤ 0.1, corresponding p-value < 0.005) [Figures 5A, Table S21] and 9 species (FDR < 0.1, corresponding p-value < 4 x 10^-4^)[Figure S10A, Table S21] were associated with circulating T_reg_ cells and TCRγδ^+^ T cells. In line with observations that supplementation of short chain fatty acids (SCFAs) augments colonic T_reg_ cell frequencies in animal models (Arpaia et al., 2013; Furusawa et al., 2013; Smith et al., 2013), the SCFA pathway “P461-PWY: hexitol fermentation to lactate, formate, ethanol and acetate” was positively associated with circulating T_reg_ cells numbers [Figure 5B]. As such, combined microbial profiling and extensive immunophenotyping illustrated that the SCFA/T_reg_ cell axis is relevant throughout adult human life. Additionally, pathways in the “Fatty Acid and Lipid Biosynthesis” superfamily, including “PWYG-321 mycolate biosynthesis”, were positively associated with circulating TCRγδ^+^ T numbers [Figure 5C], particularly in individuals in their 60s in whom TCRγδ^+^ T counts are lowest [Figure 5D]. Notably, this correlation is supported by experimental observations that TCRγδ^+^ T cells are capable of recognizing microbial lipid antigens, including mycolic acids (Ridaura et al., 2018). Together these results demonstrate that certain concepts established in animal models can be validated using human subject data. Furthermore, at the species level, the Clostridia species *Terrisporobacter othiniensis*, *Terrisporobacter glycolicus* and *Terrisporobacter glycolicus_A* were positively associated with circulating TCRγδ^+^ T cell numbers [Figure S10B]. As such, we identify both previously known microbial associations with TCRγδ^+^ T cell responses and novel associations for future mechanistic investigation.

**Figure 5.**
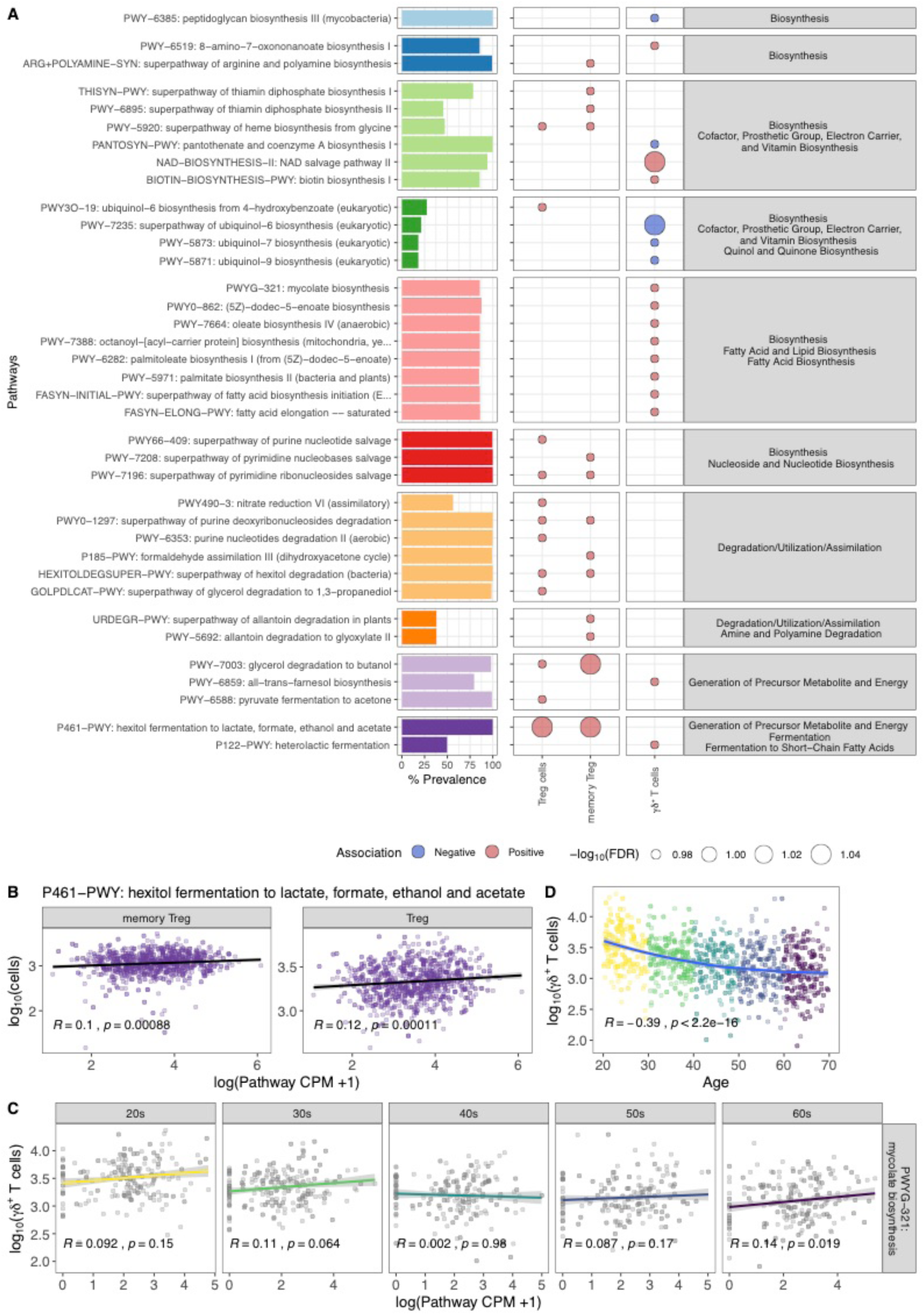
Metabolic pathways associated with circulating regulatory T (T_reg_) and γδ^+^ T cells. (A) Right bars indicate the prevalence of the pathway in the MI donors and are grouped by superpathway designation. In the center, a circle represents each significant immunophenotype x pathway interaction. Fill color indicates whether the interaction was positive/negative and size the -log10 FDR value. (B) Scatter plots show the abundance of the Short-Chain Fatty Acid pathway P461-PWY versus counts of T regulatory cells and memory cells. (C) Scatter plots show the abundance of the pathway PWYG-321 mycolate biosynthesis versus counts of γδ^+^T cells by age of the donor. (D) Scatter plot demonstrates the negative correlation between circulating γδ^+^ T cells and age of the donor. (B-D) Stats based on Spearman correlation. See also Tables S19 and S21.

We then extended our analyses to the remaining 83 immunophenotypes, in order to discover additional novel microbial/immune associations. After 45,401 tests (FDR < 0.1, corresponding p-value < 5 x 10^-5^), 20 species were associated with 17 immunophenotypes. [Figure 6A, Table S22]. Of the 24 pairs, 8 (33%) were positive associations; many of which demonstrated consistency at the phylogenetic level, for example, multiple Actinobacteriota species, including *Rothia mucilaginosa*, *Pauljensenia odontolyticus_A*, and *Pauljensenia odontolyticus_B* were positively associated with expression of the activation marker CD38 by B cells, which was previously strongly correlated with smoking (Patin et al., 2018) [Figure 6B]. Additionally, the Bacteroidota species *Bacteroides intestinalis* was negatively associated with multiple circulating T cell subsets, including both CD4^+^ and CD8^+^ central memory T cells [Figure 6A]. Further studies, requiring gastrointestinal tissue immunophenotyping, could determine whether this is due to accumulation of these cells locally within the intestinal microenvironment.

**Figure 6.**
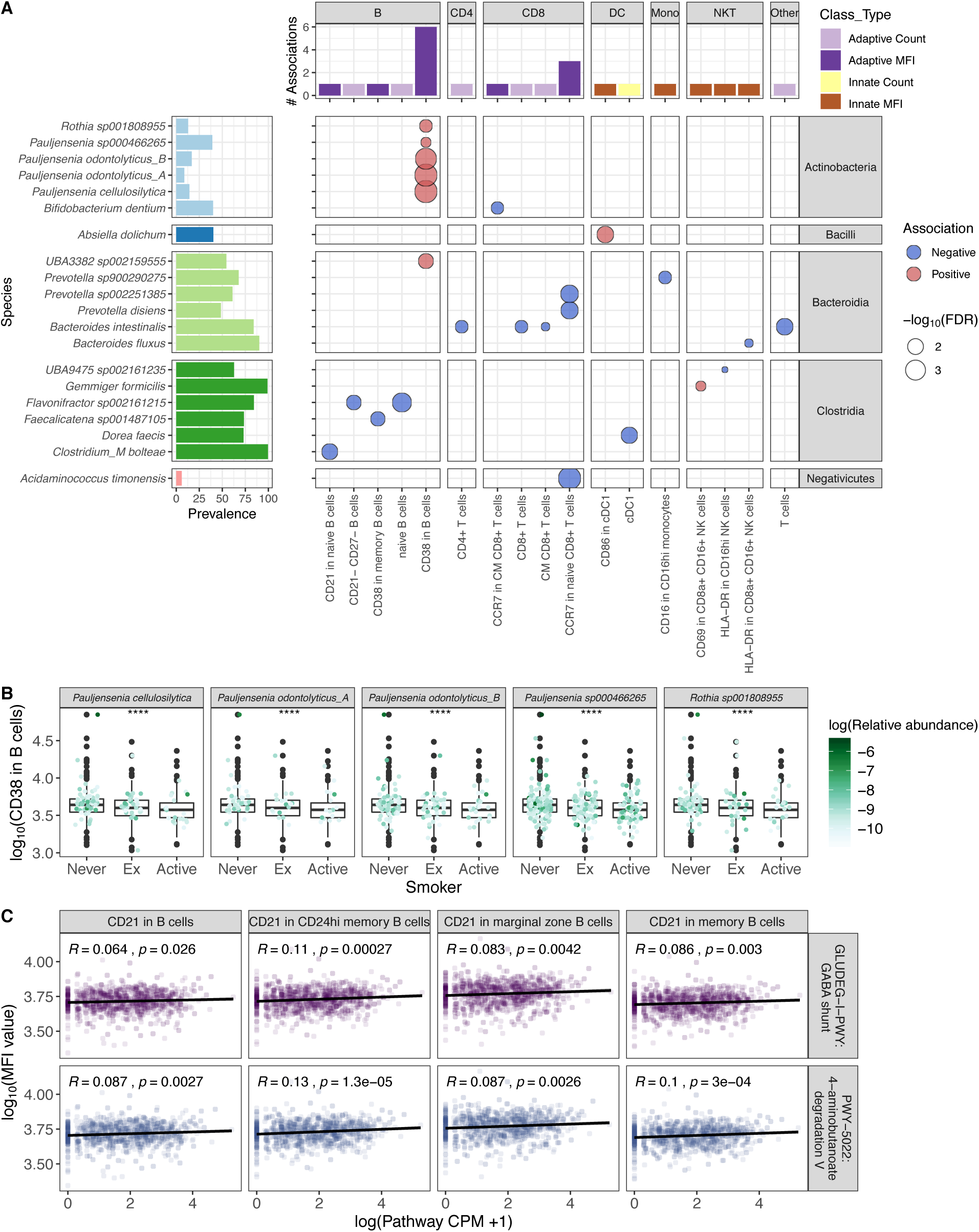
Bacterial species and metabolic pathways associated with circulating immunophenotypes. (A) Top bars indicate how many species were significantly associated with each immunophenotype. Colors indicate whether the immunophenotype relates to the adaptive or innate immune system and measures cells counts or MFI. Right bars indicate the prevalence of the species in the MI donors and are grouped by bacterial class. In the center, a circle represents each significant immunophenotype x species interaction. Fill color indicates whether the interaction was positive/negative and size the -log10 FDR value. (B) Boxplots indicate the expression of CD38 in B cells as measured by MFI by smoking status. Color corresponds to the log relative abundance of the species in the facet. **** p <= 0.0001 by Kruskal-Wallis test. (C) Scatter plots show the abundance of the pathways PWY-5022: 4-aminobutanoate degradation V and GLUDEG-I-PWY: GABA shunt pathway versus MFI for multiple B cell populations. Stats based on Spearman correlation. See also Tables S19, S20, and S22.

To further investigate the mechanisms of microbial/immune associations, after 29,050 tests (FDR < 0.1, corresponding p-value < 2.2 x 10^-5^), 3 pathways were associated with 5 immunophenotypes. [Table S22]. Notably, two pathways promoting degradation of the neuromodulator 4-aminobutanoate (GABA) were strongly associated with complement receptor 2 (CD21) expression by peripheral blood B cells [Figure 6C]. In previous association studies, the metabolic pathway “PWY-5022 4-aminobutanoate degradation V”, responsible for the breakdown of GABA into the SCFA butyrate, was identified as causally associated with increased insulin secretion following an oral glucose challenge (Sanna et al., 2019). Additionally, neuro-immunology is an emerging focus for disease pathogenesis, one that might be influenced by commensal metabolic pathways and bears further investigation (Godinho-Silva et al., 2019). Moving forward, as sample sizes increase, new microbe x immune interactions will continue to be uncovered.

## Discussion

As the number of microbial intervention trials and biomarker studies continues to grow, it is increasingly important to develop robust understanding of the gut microbiome across individuals in the steady state. In this study, we utilize the statistical power of a large cohort, the resolution afforded by deep shotgun sequencing, and an updated microbial database with genome-based taxonomy to expand on many prior microbiome observations. More specifically, we identified 28 *Prevotella* species enriched in men compared to women [Figure S6]; many of which were absent in previous databases (Truong et al., 2015) and thus not detectable in prior analyses. Given the recent literature on the strain-level variability within *Prevotella* species (De Filippis et al., 2019; Fehlner-Peach et al., 2019), particularly *P. copri*, follow-up analyses should compare if there are also strain-level differences between the sexes.

Additionally, we identified 227 species associated with age [Figure 3B], greatly expanding what was known about the effects of aging on the gut microbiome (reviewed by (An et al., 2018)). The changes seen here are particularly striking because they occur in the absence of underlying diseases or medication usage. Given the cross-sectional nature of this cohort, it is difficult to tease apart which of these associations is mediated by the variables correlated with age [Figure S4B]. For example, the interdependence of age and changes in dietary behavior could not be parsed from dependant physiological changes such as thinning of the mucosal layer or altered pH levels. To definitively understand how the gut microbiome matures with age, we will require long-term longitudinal studies. In addition, these conclusions are based on stool samples; to better understand where in the gut these changes are manifesting would require sampling along the gastrointestinal tract, as previously performed in smaller cohorts (Zmora et al., 2018).

Despite these caveats, the knowledge that many bacteria are associated with age and sex could have multiple implications for the interpretation of microbial biomarker studies. For example, several bacterial biomarkers have been reported for response to checkpoint inhibitors in non-GI cancer indications (Frankel et al., 2017; Gopalakrishnan et al., 2018; Matson et al., 2018; Routy et al., 2018). Notably, however, age, a prognostic biomarker of response and strong correlate of microbial composition, remains unaccounted for in those analyses. Additionally, these data will be valuable for designing microbial therapeutics, especially those for indications affecting older individuals. For example, interventions containing *Bifidobacterium* species may need to be dosed more frequently in individuals older than 50 years of age, in whom *Bifidobacterium* appears to colonize less effectively [Figure 3C,D, Figure S7D] . Similarly, consortia with species which demonstrated low stability in the short-term may also require additional dosing. With respect to the analysis of host immune system – microbiome associations, here we present the largest study to date to broadly associate gut bacterial profiles with circulating immune cells. We used this information to successfully validate results from animal studies in human subject data, including the association of SCFAs with T regulatory cells as well as microbial fatty acids and TCRγδ^+^ T cells. While this dataset is well powered for targeted questions, additional samples will support the discovery of novel associations with smaller effect sizes. Finally, beyond the findings in this manuscript, the rich metadata and 1,000 plus deep shotgun metagenomic samples provided here will be a valuable resource upon which future microbiome studies can test and build new computational tools as well as generate and test new hypotheses.

## Acknowledgements

We would like to thank all the donors for their participation to the study. We thank J. Segre and F. Tamburini for critical reading of the manuscript and helpful comments.

Author: The Milieu Intérieur Consortium^†^.

† The Milieu Intérieur Consortium¶ is composed of the following team leaders: Laurent Abel (Hôpital Necker), Andres Alcover, Hugues Aschard, Kalla Astrom (Lund University), Philippe Bousso, Pierre Bruhns, Ana Cumano, Caroline Demangel, Ludovic Deriano, James Di Santo, Françoise Dromer, Gérard Eberl, Jost Enninga, Jacques Fellay (EPFL, Lausanne), Ivo Gomperts-Boneca, Milena Hasan, Serge Hercberg (Université Paris 13, Paris), Olivier Lantz (Institut Curie, Paris), Hugo Mouquet, Etienne Patin, Sandra Pellegrini, Stanislas Pol (Hôpital Côchin, Paris), Antonio Rausell (INSERM UMR 1163 – Institut Imagine, Paris), Lars Rogge, Anavaj Sakuntabhai, Olivier Schwartz, Benno Schwikowski, Spencer Shorte, Frédéric Tangy, Antoine Toubert (Hôpital Saint-Louis, Paris), Mathilde Trouvier (Université Paris 13, Paris), Marie-Noëlle Ungeheuer, Darragh Duffy§, Matthew L. Albert (In Sitro)§, Lluis Quintana-Murci§,

¶ unless otherwise indicated, partners are located at Institut Pasteur, Paris, France

§ co-coordinators of the Milieu Intérieur Consortium

Additional information can be found at: http://www.pasteur.fr/labex/milieu-interieur

This work benefited from support of the French government’s Invest in the Future programme. This programme is managed by the Agence Nationale de la Recherche, reference ANR-10-LABX-69-01.

## Author Contributions

Conceptualization, A.L.B., O.J.H., D.D., M.L.A, L.Q-M.; Methodology, A.L.B., M.L., J.B.; Software, A.L.B., M.L.; Formal Analysis, A.L.B., S.L., M.L., K.E.F.; Investigation, B.C.; Resources, D.D., M.L.A., L.Q-M,;Data Curation, A.L.B., J.B.; Writing – Original Draft, A.L.B., O.J.H.; Writing – Review & Editing, D.D, S.L., E.P., D.N., M.L., K.E.F., J.B., M.L.A.; Visualizations, A.L.B., M.L.; Supervision, D.N., D.D., M.L.A.; Project Administration, D.N., D.D.; Funding Acquisition, D.D., M.L.A., L.Q-M.

## Methods

### EXPERIMENTAL MODEL AND SUBJECT DETAILS

#### The Milieu Intérieur cohort

The 1000 healthy donors of the Milieu Intérieur cohort were recruited by BioTrial in the suburban Rennes area (Ille-et-Vilaine, Bretagne, France). The cohort included 500 men and 500 women, and 200 individuals from each decade of life, between 20 and 69 years of age. Participants were selected based on stringent inclusion and exclusion criteria, detailed elsewhere (Thomas et al., 2015). Donor BMI was restricted to ≥ 18.5 and ≤ 32 kg/m^2^. Briefly, they had no evidence of any severe/chronic/recurrent pathological conditions. Primary exclusion criteria were seropositivity for human immunodeficiency virus or hepatitis C virus, travel to (sub-) tropical countries within the previous 6 months, recent vaccine administration, and alcohol abuse. Subjects were also excluded if they took nasal, intestinal, or respiratory antibiotics or antiseptics any time in the 3 months preceding enrollment. Additionally, anyone following a doctor or dietician prescribed diet for medical reasons (e.g. calorie-controlled diet in overweight patients) and volunteers with food intolerance or allergy were excluded. To avoid the influence of hormonal fluctuations in women during the peri-menopausal phase, only pre- or post-menopausal women were included. To minimize the influence of population substructure, the study was restricted to individuals of self-reported Metropolitan French origin for three generations (i.e., with parents and grandparents born in continental France).

#### Demographic, environmental, dietary, and clinical variables

Multiple demographic, environmental, and clinical variables were collected for each of the donors in an electronic case report form (Thomas et al., 2015). For example, donors were asked about their family medical history, smoking habits, sleeping habits, and infection and vaccination history. Additionally, donors completed a food-frequency questionnaire (FFQ) administered by trained investigators and comprising 19 food groups [Table S5]. Participants estimated their “usual consumption” selecting from 6 intake frequencies ranging from “Twice per day or more” to “Never” (except for alcohol, which offered 5 intake frequencies ranging from “Every day” to “Never”). Investigators administering the FFQ invited participants to declare their “usual” diet, rather than focusing on their latest dietary consumption. The detailed FFQ is available in (Partula et al., 2019). For clinical chemistry, hematologic, and serologic assessments, 20mL of blood was collected from each donor and analyzed at the certified *Laboratoire de biologie m*é*dicale, Centre Eugene Marquis* (Rennes, France).

After manual curation and removal of variables that were (i) variable in less than 5% of participants (ii) missing in more than 25% of donors (iii) correlated with another variable Spearman Rho > -0.6 or < 0.6, 154 metadata variables were considered for future associations. In the case of correlated variables [Table S5,6], the variable with fewer missing values was prioritized and kept, while the other variable was removed. When the pair had equivalent numbers of missing values, one from the pair was randomly selected. Notably, circulating levels of creatinine were so strongly correlated with sex (Spearman Rho = 0.72, p-value 3.5 e-115), this variable was excluded from the 154.

#### Ethics Statement

The clinical study was approved by the Comité de Protection des Personnes - Ouest 6 on June 13, 2012, and by the French Agence Nationale de Sécurité du Médicament on June 22, 2012, and was performed in accordance with the Declaration of Helsinki. The study was sponsored by the Institut Pasteur (Pasteur ID-RCB Number 2012-A00238-35) and conducted as a single-center study without any investigational product. The original protocol is registered under ClinicalTrials.gov (study number NCT01699893). Informed consent was obtained from the participants after the nature and possible consequences of the studies were explained. The samples and data used in this study were formally established as the Milieu Interieur biocollection (NCT03905993), with approvals by the Comité de Protection des Personnes – Sud Méditerranée and the Commission nationale de l’informatique et des libertés (CNIL) on April 11, 2018.

### METHOD DETAILS

#### Fecal DNA extraction and shotgun metagenomic sequencing

Subjects were asked to produce stool samples at their home within 24 hours before the scheduled visits (V1, V2). Stool specimens were collected in a double-lined sealable bag containing a GENbag Anaer atmosphere generator (Aerocult, Biomerieux) to maintain anaerobic conditions. Upon reception at the clinical site, fresh samples were aliquoted into cryotubes and stored at -80 °C.

Stool aliquots were shipped to the CRO Diversigen for DNA extraction and shotgun metagenomic sequencing. At Diversigen, genomic DNA was extracted using PowerMag Soil DNA Isolation Kit (Qiagen - MO BIO Laboratories, Catalog No. 27100). Libraries were prepared using Beckman robotic workstations (Biomek FX and FXp models) in batches of 96 samples. DNA (10 ng to 500 ng) was sheared into fragments of approximately 300-400 bp in a Covaris E210 system (96 well format, Covaris, Inc. Woburn, MA) followed by purification of the fragmented DNA using AMPure XP beads. DNA end repair, 3’-adenylation, ligation to Illumina multiplexing PE adaptors, and ligation-mediated PCR (LM-PCR) was all completed using automated processes. To amplify high GC and low AT rich regions at greater efficiency, KAPA HiFi polymerase (KAPA Biosystems Inc.) was used for PCR amplification (6-10 cycles). Fragment Analyzer (Advanced Analytical Technologies, Inc) electrophoresis system was used for library quantification and size estimation. Prepared libraries were then pooled and sequenced on an Illumina HiSeq 2500.

In the end, we obtained 21 trillion raw paired-end reads from 1,363 samples from 946 of the donors. On average per sample, there were 2.4 Gbp, 15.5 million reads with 358 bp insert size. To process the reads, Illumina TruSeq adapters were trimmed with Trimmomatic v0.36 (Bolger et al., 2014); low quality and low complexity reads were removed with prinseq-lite 0.20.4 (Schmieder and Edwards, 2011); and Bowtie2 v2.1.0 (Langmead and Salzberg, 2012) was used to remove reads mapping to PhiX or the PacBio human genome (parameters specified in Fig S1C). After processing, there were on average 14.2 ± 2.8 million reads per sample [Fig S1D, E]. Out of an initial 1,000 recruited donors, 44 donors were excluded from this analysis due to a lack of consent for sharing their data outside of the MI consortium. An additional 10 donors were excluded because of technical issues in the extraction and the sequencing steps (e.g., due to low DNA extraction yield) resulting in a sample size of 946 donors.

### QUANTIFICATION AND STATISTICAL ANALYSIS

#### Building the Kraken-GTDB database

To build the Kraken-GTDB database, first the following files were downloaded ftp://ftp.ncbi.nlm.nih.gov/genomes/refseq/bacteria/assembly_summary.txt and https://data.ace.uq.edu.au/public/gtdb/data/releases/release89/89.0/bac120_taxonomy_r89.tsv (Parks et al., 2018) on 2019-06-25. These files were merged based on accession number and only those genomes present in both databases were considered, aka RefSeq genomes with a GTDB taxonomy. To avoid biasing the database towards those species with large numbers of genomes [Figure S2A], while balancing the added information provided by additional isolates per species, we selected 5 genomes per GTDB species to include in our database. Genomes were first ordered by their assembly quality, i.e. reference genome, representative genome, Complete Genome, Chromosome, Contig, Scaffold and then randomly selected. Based on this criteria, 23,505 genomes representing 13,446 unique bacterial species were downloaded and formatted into a Kraken2 database (Wood and Salzberg, 2014). To incorporate the GTDB taxonomy into the Kraken2 database, files mimicking the “NCBI-like” taxonomy files from ftp://ftp.ncbi.nih.gov/pub/taxonomy/new_taxdump/new_taxdump.zip were created for names.dmp, complete_names.dmp, nodes.dmp, and accession2taxid. A matching Bracken database was then generated with bracken-build -k 35 -l 126 (Lu et al., 2017).

#### Metagenomic data analysis

First, putative reagent contaminants identified by species co-correlation analysis were filtered using Kraken2’s *-unclassified-out* option and a custom database of contaminant genomes [Figure S11, Table S23]. Using our custom GTDB-Kraken database, Kraken2 v2.0.6-beta (Wood and Salzberg, 2014) and Bracken v2.5 (Lu et al., 2017) were run on the 1,363 quality-controlled samples (parameters specified in Figure S1F) to generate bacterial profiles. With the exception of Figure 4, all analyzes were based on bacterial profiles from the Visit 1 samples. In the case no Visit 1 sample was available, the sample from Visit 2 was used.

To complement results from the MI donors, this pipeline was run on an additional 1,159 samples from 851 donors downloaded from ENA: PRJEB11532 (Zeevi et al., 2015). Notably, the reagent contaminants identified in the MI donors were nearly absent in the Zeevi samples, so no reagent filtering was performed. For these samples, metadata (Sex, Age, BMI) was obtained by emailing the authors. No timepoint was provided, so an average of the microbial profiles across samples of the same donor were used in further analyzes.

To go beyond taxa-based calls, HUMANn2 v0.11.2 (Franzosa et al., 2018) with default parameters including Uniref90 was run on all MI samples where the forward and reverse reads were concatenated into a single file. Raw output values were converted from *rpks* to *cpms* with *humann2_renorm_table*. Outputs of all samples were joined into a single merged table with the *humann2_join_tables* function. Using *humann2_regroup_table*, individual gene families were regrouped with multiple different databases including COG, GO, KEGG, and MetaCyc. Despite varying magnitudes of richness, at a high level, results were consistent across the different databases [Figure S12]. Ultimately, MetaCyc pathways (Caspi et al., 2018) were selected for the associations because of the additional steps implemented in HUMANn2 to check for completeness of the pathways.

#### Statistical Analysis

All associations and statistical tests were performed in R v3.6.1 (R Core Team, 2019), documented via rmarkdown documents (Allaire et al., 2019), and compiled with knitr (Xie, 2019). Within R, tables were manipulated with functions of the dplyr package (Wickham et al., 2019). The majority of figures were rendered with ggplot2 (Wickham, 2016) and arranged with cowplot (Wilke, 2019). Colors were selected with the help of RColorBrewer (Neuwirth, 2014) and viridis (Garnier, 2018). The cladogram in Figure 3B was generated with GraPhlAn (Asnicar et al., 2015). Correlation plots in Figure S4, S7C were generated with ggcorrplot (Kassambara, 2019). Supplemental tables were generated with Openxlsx (Walker, 2019). When comparing values between two or more groups, Wilcoxon-Rank Sum tests were used.

Bacterial α- and β-diversity measures including Shannon and Bray-Curtis were calculated using the R package vegan (Oksanen et al., 2019). To identify which of the 154 metadata variables were significantly associated with Bray-Curtis β-diversity, we used the adonis function in vegan to run permutational analysis of variance (PERMANOVA) tests with 999 permutations. To identify bacterial species and pathways differen abundant between males and females, we used DESeq2 (Love et al., 2014). To identify bacterial species and pathways differentially prevalent between males and females, we used prop.test in R. Bacterial taxa and pathways associated with age were identified using Spearman correlations.

To determine the stability of a donor’s bacterial species and pathways between Visit 1 and Visit 2, the Bray-Curtis distance was calculated and subtracted from 1. Individuals with a stability of 1 had samples that were identical across timepoints, while a stability of 0 meant the samples were nothing alike. Similarly, 1 - the Jaccard index was used to determine pathway stability based on presence/absence. To identify the phyla associated with stability, Spearman correlation coefficients were calculated. We then used generalized linear mixed models (GLMMs) fit with the R package glmmTMB (v.0.2.3)(Mollie E. Brooks et al., 2017) with a beta response to identify which metadata factors were associated with intra−individual stability. To calculate stability of individual features (species and pathways), we again applied the Bray-Curtis metric but this time compared the relative abundance of a single feature across all donors at Visit 1 and Visit 2.

Bacterial profiles (species and pathways) were associated with circulating immunophenotypes with multivariate linear models, akin to the procedure in (Patin et al., 2018). To reduce the multiple testing burden, of the 166 immunophenotypes originally published (Patin et al., 2018), we prioritized the 40 cell counts that had a mean count greater than 1,000 and their 46 associated MFI intensities [Figures S8,9, Tables S19,20]. Fifty of these immunophenotypes described adaptive immune system, while 36 were for innate cells. With multivariate linear modeling using the lmer function from lmerTest R package (Kuznetsova et al., 2017), correcting for age (spline, df = 3), sex, CMV status, smoking, genetics, and batch which were previously associated with immune profiles (Patin et al., 2018), 547 bacterial species (prevalence >5%, average abundance > 0.01%, TSS normalized) were associated with the 86 immunophenotypes for 47,042 tests. Similarly, 350 metabolic pathways (prevalence > 20%) were correlated with the 86 immunophenotypes with the same linear models for 29,050 tests. More specifically, the formula utilized for each combination was

immunophenotype ∼ pathway (or species) + SNPs + ns(Age, df = 3) + Sex + Smoking + TABAC.T1 +AncestryPC1 + AncestryPC2 + HourOfSampling + (1 | DateOfSampling),

where *immunophenotypes* and *species* are log-transformed, if contains 0, then 1 unit is added before log-transformation; the *pathway* is the log(CPM + 0.6884654), where 0.6884654 is the smallest pathway CPM value.

For all statistical tests, p-values were corrected with the R function p.adjust using the Benjamini & Hochberg (FDR) method.

### DATA AND SOFTWARE AVAILABILITY

#### Data Availability

Sequence data will have been deposited in the European Genome-Phenome Archive under accession code EGAXXX prior to acceptance. Donor metadata and code utilized in this paper will also be available.

## Supplemental Figures

**Figure S1.**
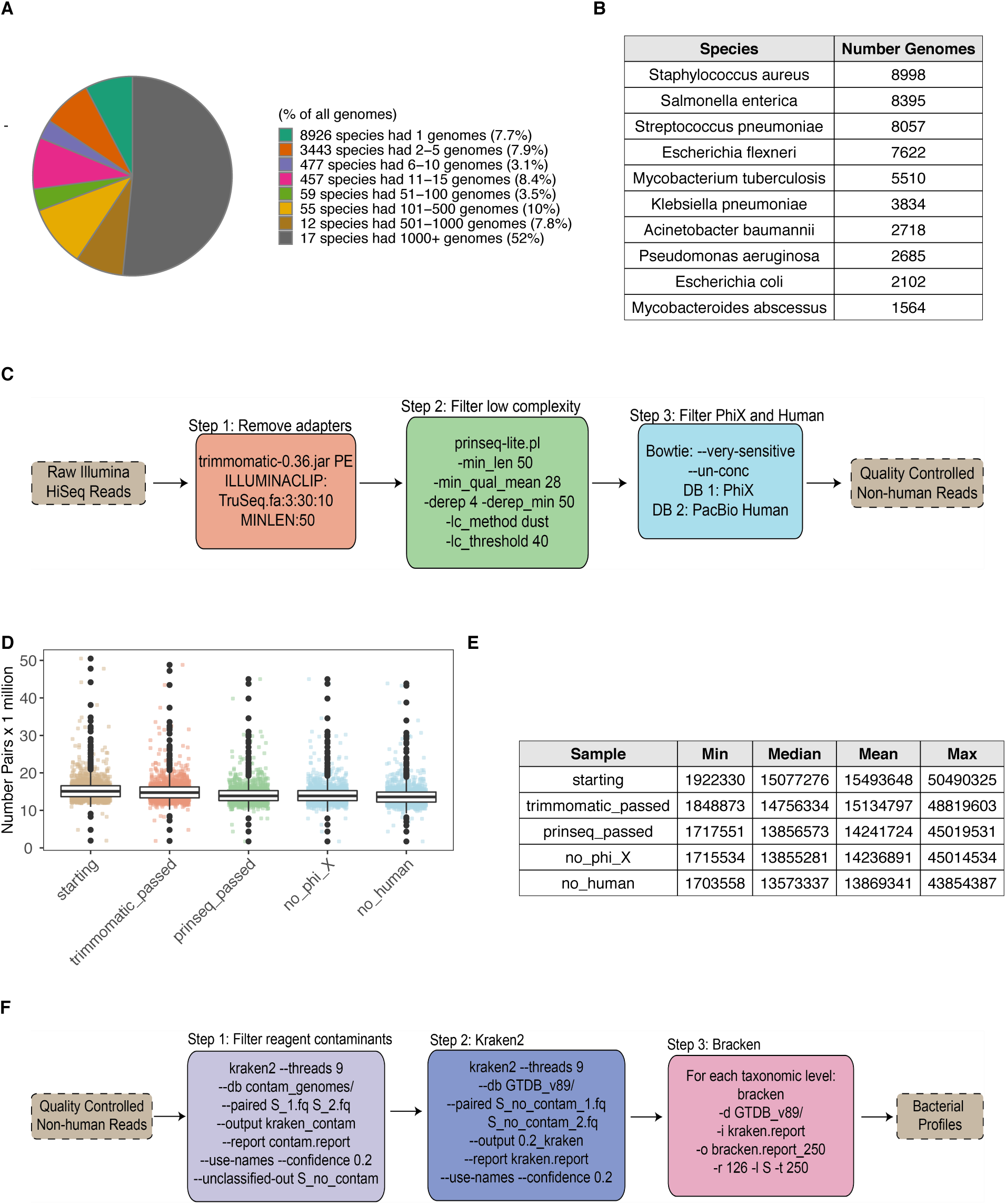
Quality control and Kraken analysis pipeline, Related to Figure 1. (A) Pie chart indicating the distribution of genomes in RefSeq (as of 2019-06-25) to different species, based on GTDB taxonomy. 29 species represent over 60% of the reference genomes. (B) Table of the 10 species with the greatest number of genomes in RefSeq. (C) Steps taken to quality control shotgun metagenomic reads. Additional details can be found in the Methods Section. (D) Barplots show the number of paired-end reads remaining after each filtration step in (C). Each point corresponds to 1 sample (n = 1363). Colors correspond to steps in (C). (E) Summary table for the number of paired-end reads in (D). (F) Steps taken to run Kraken2 and Bracken on the quality-controlled reads including removal of putative reagent contaminants. Additional details can be found in the Methods Section. See also Table S2.

**Figure S2.**
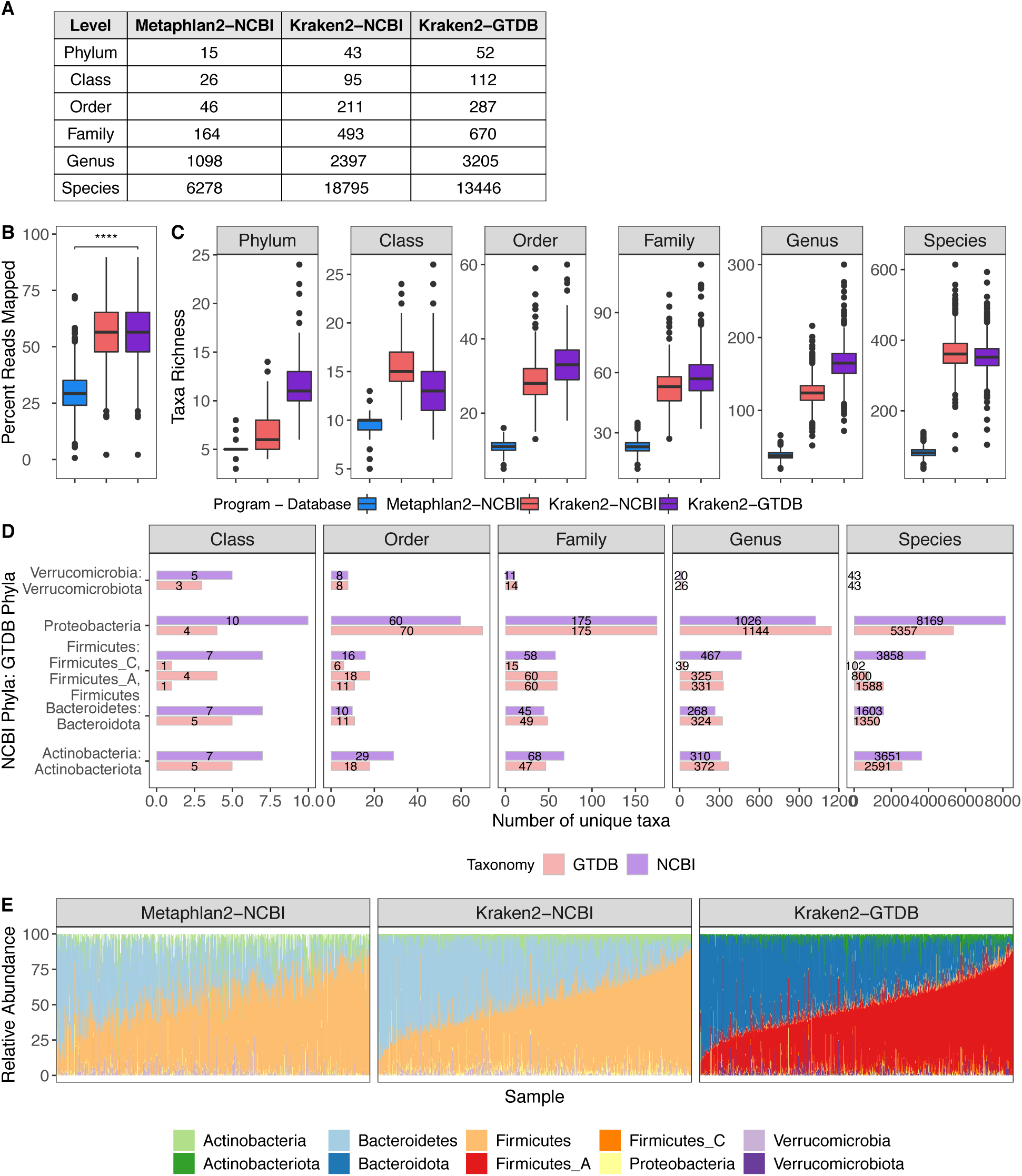
Comparison of all Program-Database combinations tested, Related to Figure 1. (A) Table indicates the richness across taxonomic levels of the different databases tested. The NCBI and Genome Tree Database (GTDB) databases used by Kraken2 include the same 23505 RefSeq genomes with taxonomies assigned either by NCBI or the GTDB respectively. GTDB taxonomic assignments often lead to collapsing or splitting of those taxa originally in NCBI; e.g.Firmicutes was split into Firmicutes, Firmicutes_A, Firmicutes_B, Firmicutes_C, etc. (B) Boxplots compare the percent of shotgun metagenomic reads mapped by the different program-database combinations. Percentage of mapped reads for Metaphlan2 was based on its estimated number of reads from the clade. **** p < 0.0001 by Wilcoxon-Rank Sum. (C) Boxplots compare the richness of taxa identified per sample with the different program-database combinations tested. All pairwise comparisons had p < 0.0001 by Wilcoxon-Rank Sum. (D) Bars indicate the number of unique taxa present in the NCBI (purple) and GTDB (red) databases for each of the top phyla across taxonomic levels. (E) Relative abundance of the top phyla across different program-database combinations. Each bar corresponds to 1 sample. Samples are ordered based on the relative abundance of Firmicutes_A found by Kraken2-GTDB.

**Figure S3.**
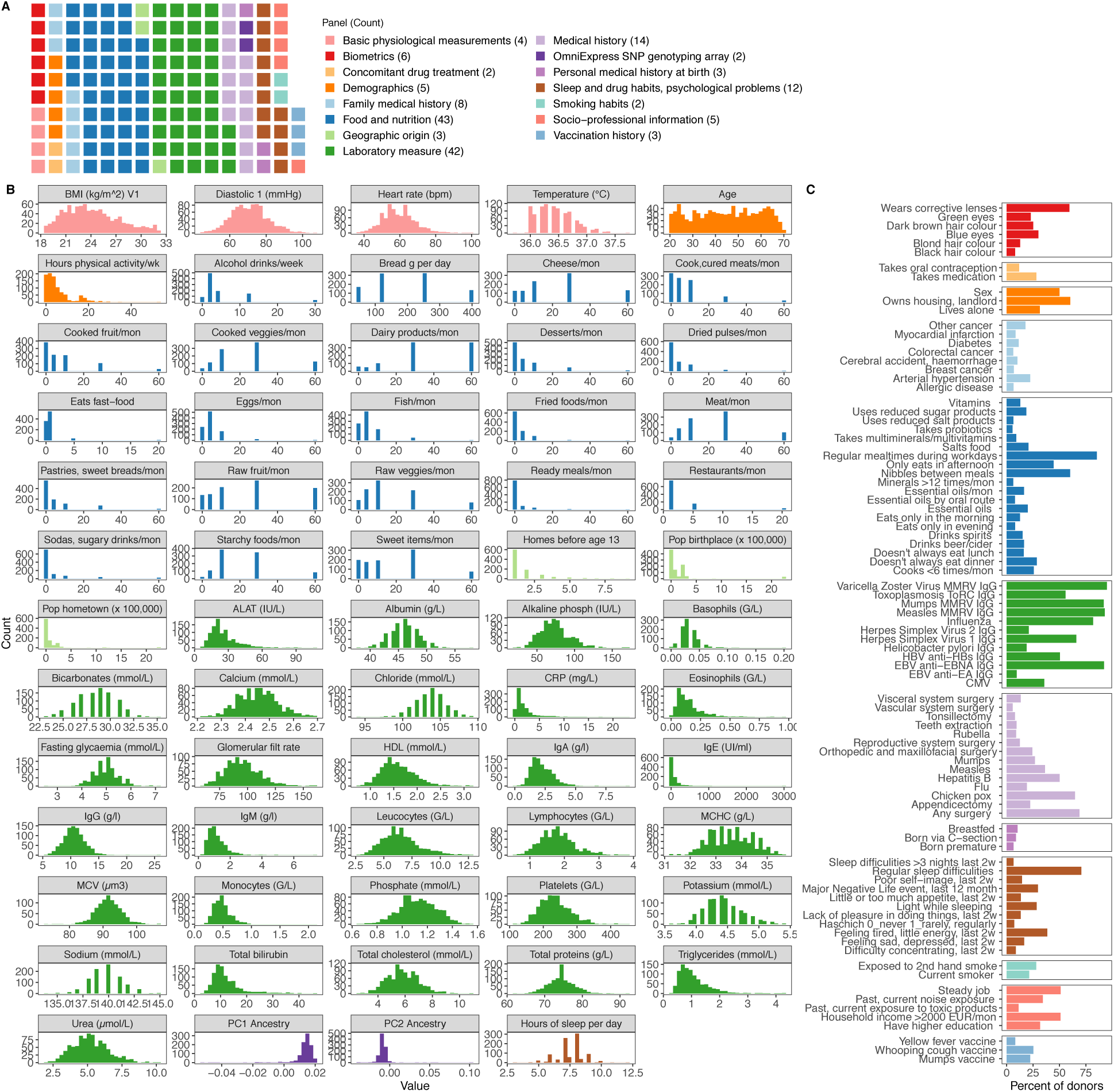
154 variables were associated with bacterial profiles, Related to Figure 1. (A) Distribution of variables across broad categories. (B) Distribution of 64 continuous variables across donors. Colors correspond to those in A. (C) Percent of donors for which each of the 90 binary variables was True. Colors correspond to those in A. 2w = 2 weeks, ALAT = Alanine Aminotransferase, CMV = Cytomegalovirus, CRP = C-Reactive Protein, EBV = Epstein-Barr virus, G/L = billion of cells per liter, HBV = Hepatitis B virus, HDL = high-density lipoproteins, Ig = Immunoglobulin, MCV = Mean corpuscular volume, MCHC = Mean corp hemoglobin concentration, mon = month, PC = Principal Component of genetics SNP array See also Tables S5 and S6.

**Figure S4.**
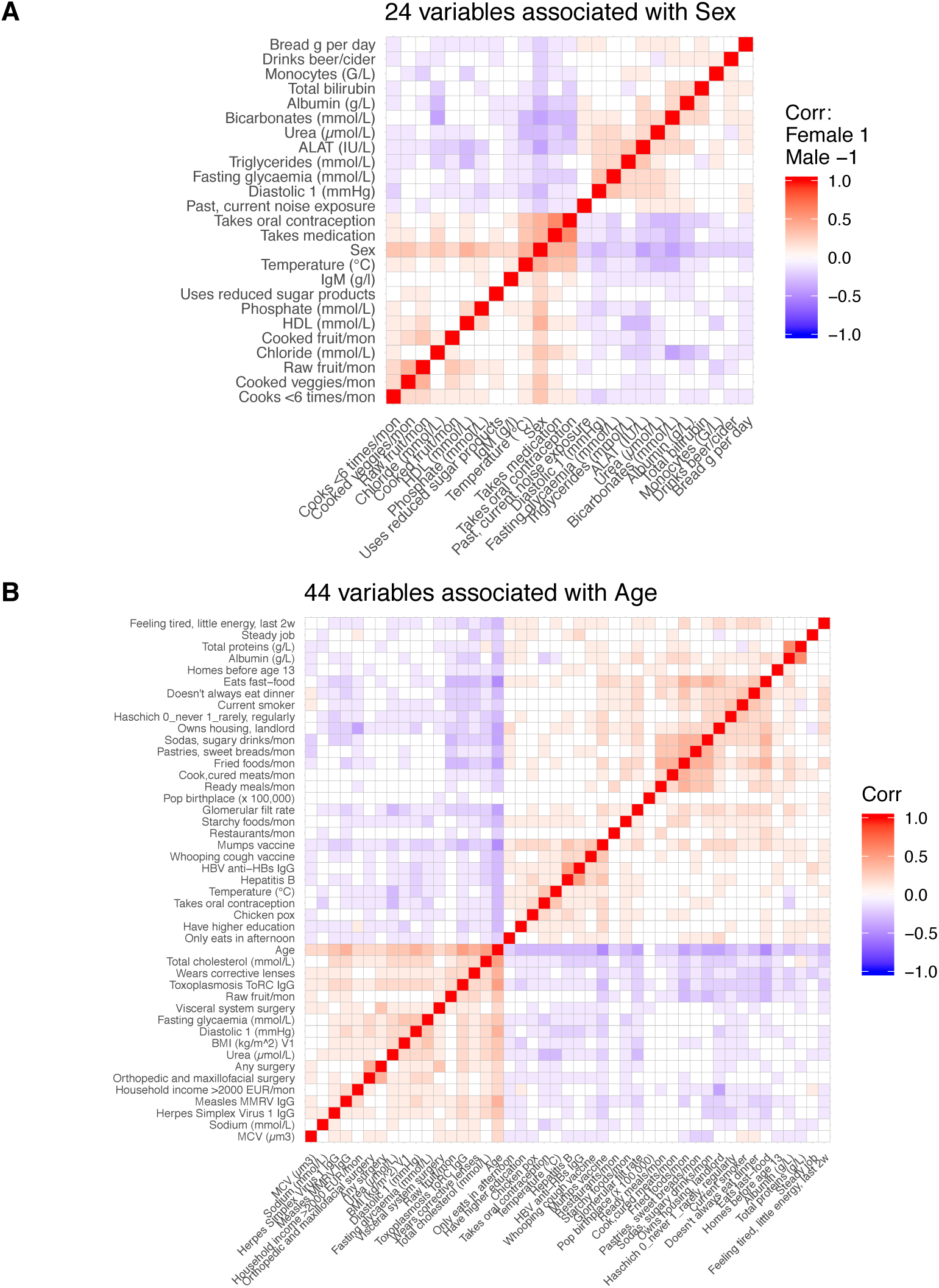
Metadata variables co-correlated with sex and age, Related to Figure 1. (A) 24 variables were associated with sex with Spearman Rho > 0.2 or < -0.2. (B) 44 variables were associated with age with Spearman Rho > 0.2. or < -0.2. See also Table S6.

**Figure S5.**
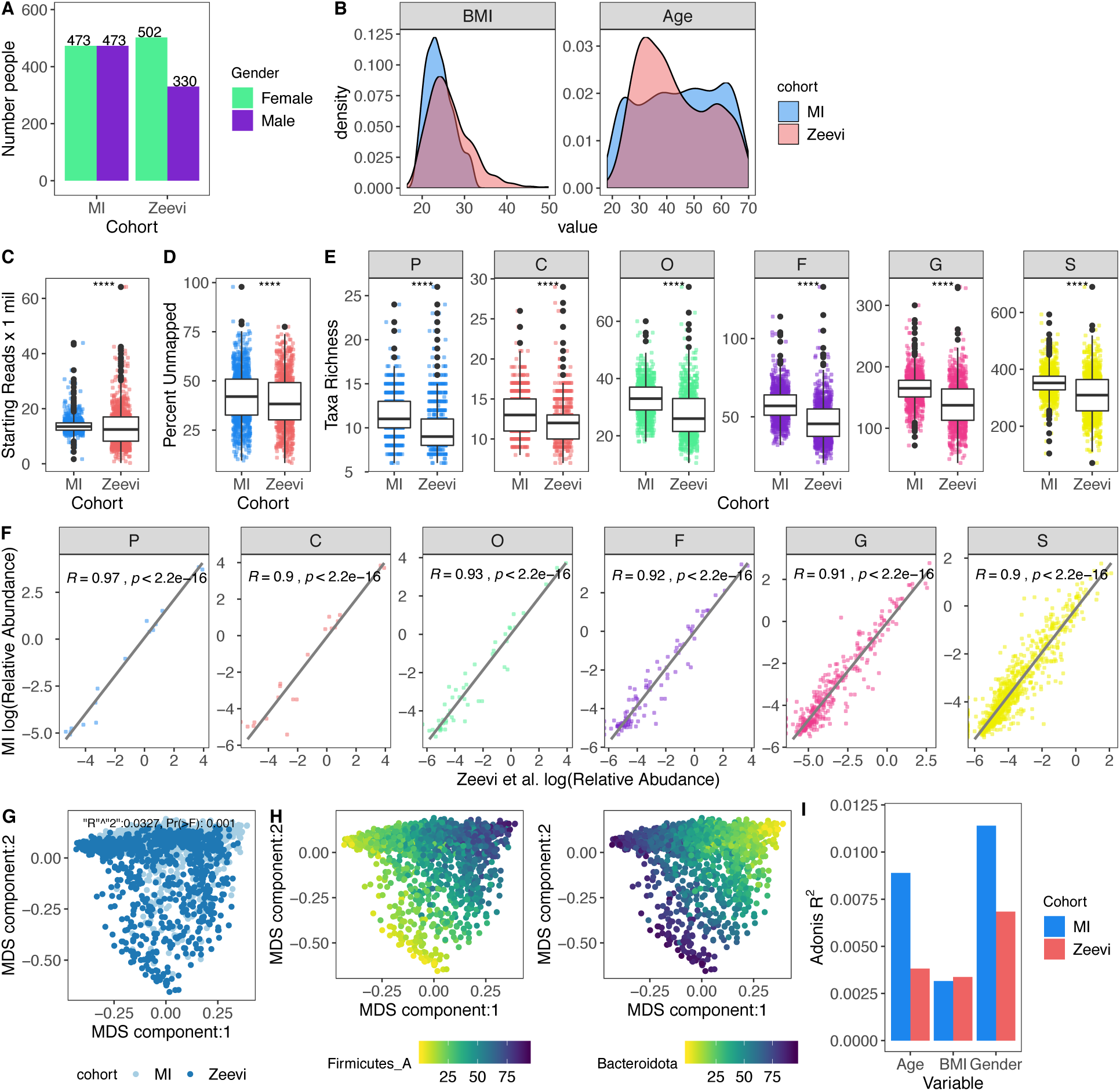
Microbial profiles of MI donors in comparison to those from Zeevi et al., Related to Figure 1. (A) Bars indicate the number of females and males in both cohorts. (B) Density plots show distribution of Body Mass Index (BMI) and Age in both cohorts. (C-E) **** p < 0.0001 by Wilcoxon-Rank Sum. (C) Bar plots compare sequencing depth across cohorts. (D) Barplots show percent of reads unmapped after the Kraken2-GTDB pipeline. (E) Boxplots show richness across taxonomic levels. Each dot corresponds to 1 donor. (F) Association of relative abundances of each taxon across taxonomic levels with Spearman correlation. Each dot corresponds to 1 taxon. Only taxa present in >5% of either cohort were considered. Log relative abundances are shown. (G) Multidimensional scaling (MDS) ordination plot of all samples in both the MI and Zeevi cohorts. Each dot corresponds to 1 donor and is colored by the cohort. R^2^ indicates the amount of interindividual variation (calculated with Bray Curtis) explained by the cohort and was calculated with the PERMANOVA test adonis. (H) MDS ordination plot of samples in both the MI and Zeevi cohorts. Each dot corresponds to 1 donor and is colored by the relative abundance of the phyla. (I) Bars indicate the amount of interindividual variation (calculated with Bray Curtis) explained by each of the variables in each of the cohorts. See also Table S8.

**Figure S6.**
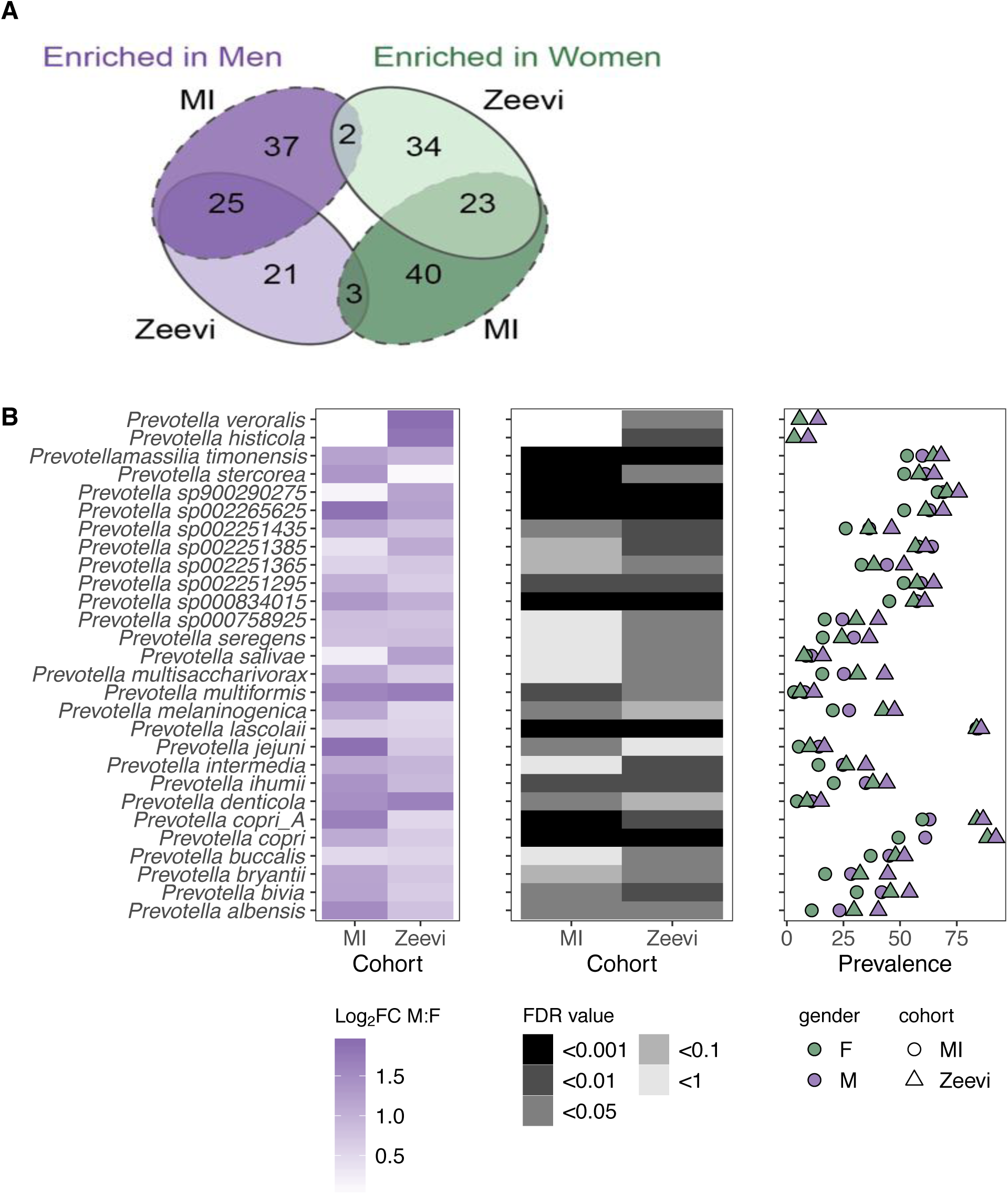
Associations of taxa with sex, particularly Prevotella, were consistent in the Zeevi cohort, Related to Figure 2. (A) Venn diagram comparing all the bacterial species statistically significantly (FDR < 0.05) associated with sex in the MI and Zeevi cohorts. (B) 28 *Prevotella* species were more abundant in males consistently across cohorts. Left panel shows the log2 fold change (Log2FC) of species in males versus females. Middle panel indicates the FDR value. Notably, even when a species was significant in only one cohort the direction was consistent in the other. Right panel shows the prevalence of each species. Color indicates the sex, while shape indicates the cohort. See also Tables S9 and S10.

**Figure S7.**
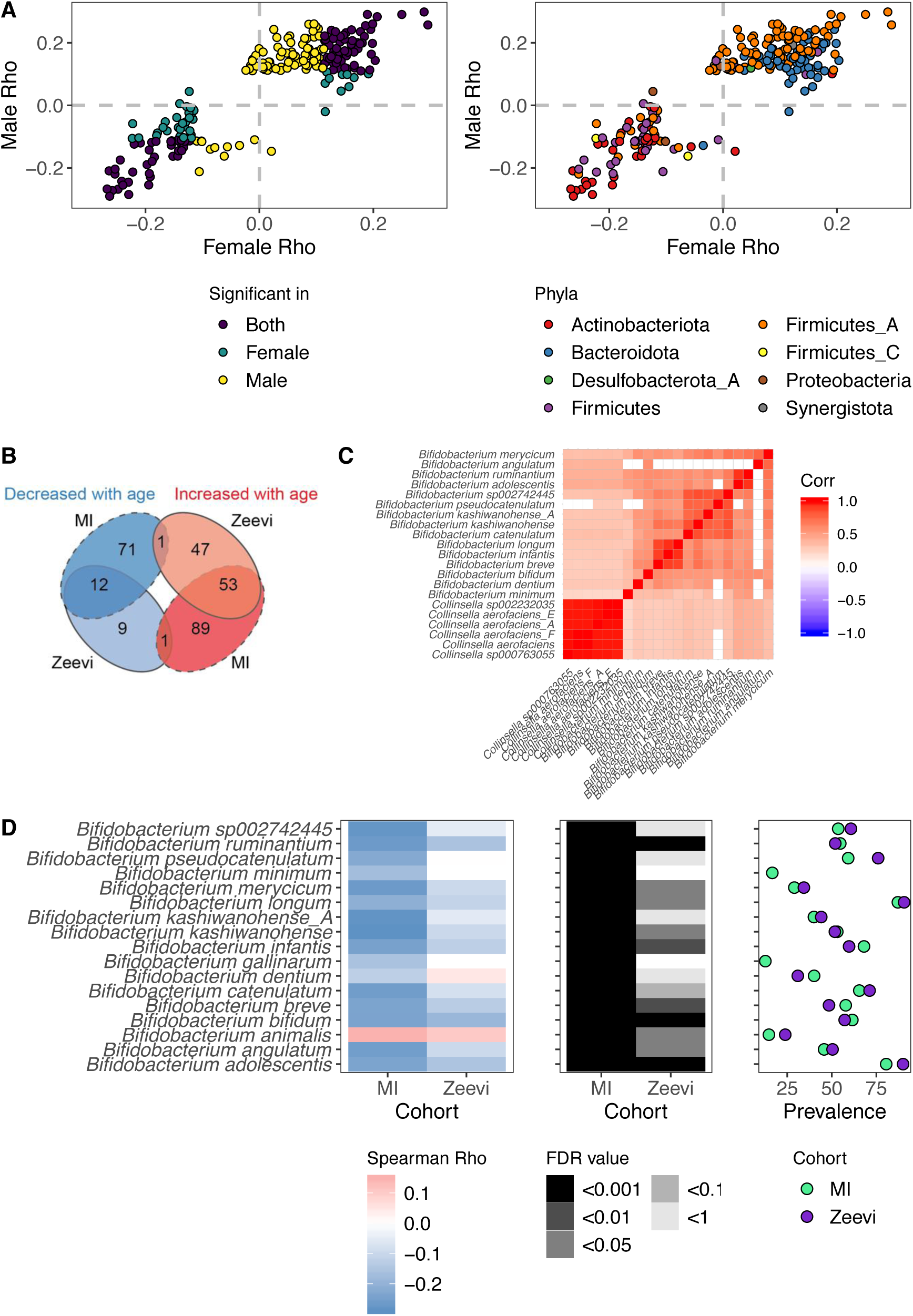
Associations of taxa with age, particularly Bifidobacterium, were consistent across sexes and in the Zeevi cohort, Related to Figure 3. (A) Scatter plots compare the Spearman rho values of bacteria ∼ age for males and females, 1 point = 1 species. In the left plot, points are colored based on whether the association was significant (FDR < 0.05) in males, females, or both. In the right plot, points are colored based on the phyla designation of the species. (B) Venn diagram comparing the bacterial species statistically (FDR < 0.05) associated with age in the MI and Zeevi cohorts. (C) Correlation plot of *Bifidobacteria* species (prevalence > 30%) and their co-correlated *Collinsella* species (Spearman Rho > 0.4) (D) 17 *Bifidobacterium* species were associated with age across cohorts. Left panel shows the association of species with age (Spearman Rho). Middle panel indicates the FDR value. Notably, even when a species was significant in only one cohort the trend was consistent in the other. Right panel shows the prevalence of each species. Color indicates the cohort. See also Tables S12 and S13.

**Figure S8.**
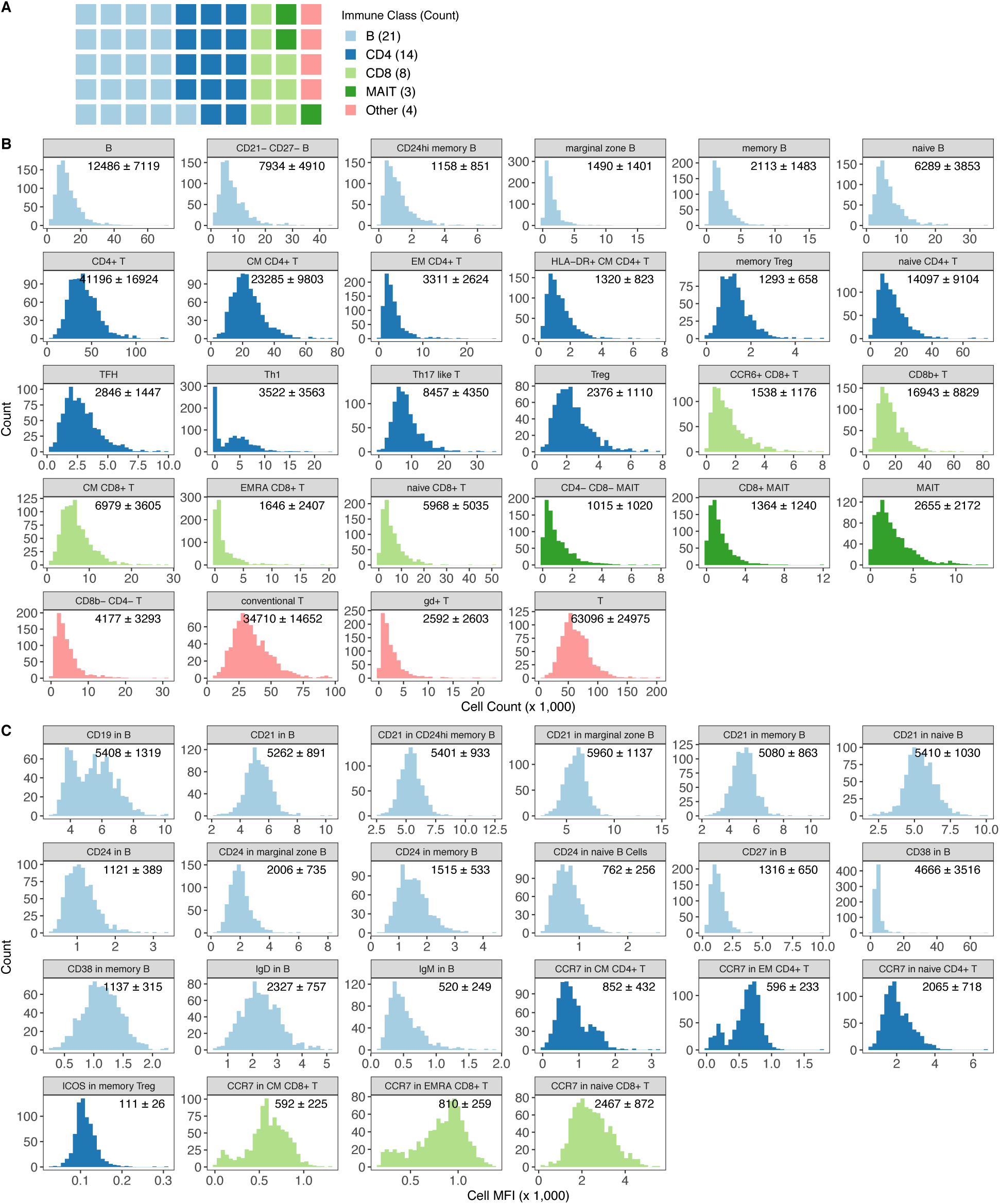
50 circulating adaptive immunophenotypes were associated with bacterial profiles, Related to Figures 5 and 6. (A) Distribution of adaptive immunophenotypes across cell type. (B) Distribution of 28 circulating immune cell counts across donors. Colors correspond to those in A. Numbers are mean cell count ± standard deviation. (C) Distribution of 22 Mean Fluorescence Intensities (MFIs) across donors. Colors correspond to those in A. Numbers are mean value ± standard deviation. See also Tables 19 and 20.

**Figure S9.**
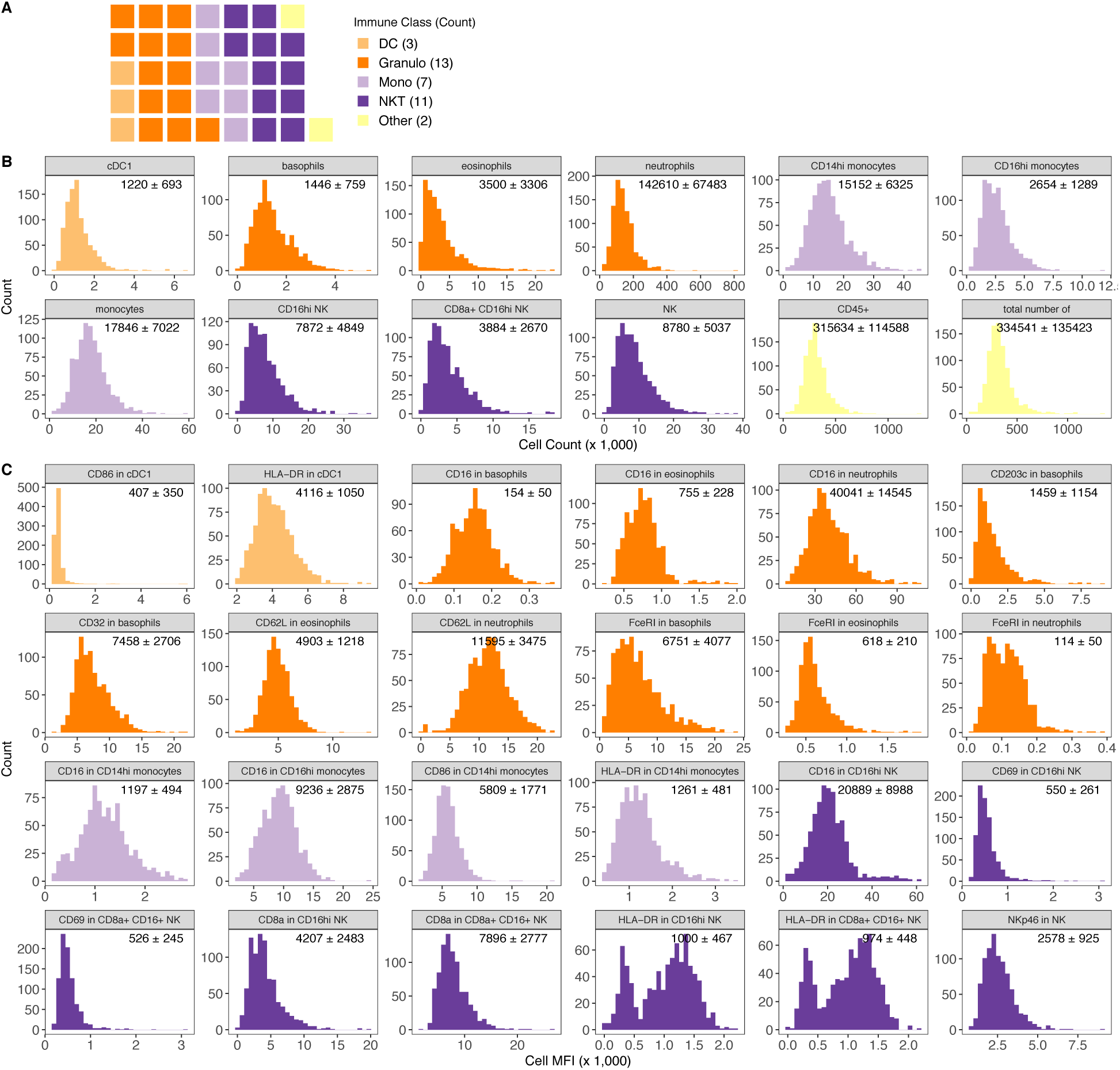
36 circulating innate immunophenotypes were associated with bacterial profiles, Related to Figures 6. (A) Distribution of innate immunophenotypes across cell type. (B) Distribution of 12 circulating immune cell counts across donors. Colors correspond to those in A. Numbers are mean cell count ± standard deviation. (C) Distribution of 24 Mean Fluorescence Intensities (MFIs) across donors. Colors correspond to those in A. Numbers are mean value ± standard deviation. See also Tables S19 and S20.

**Figure S10.**
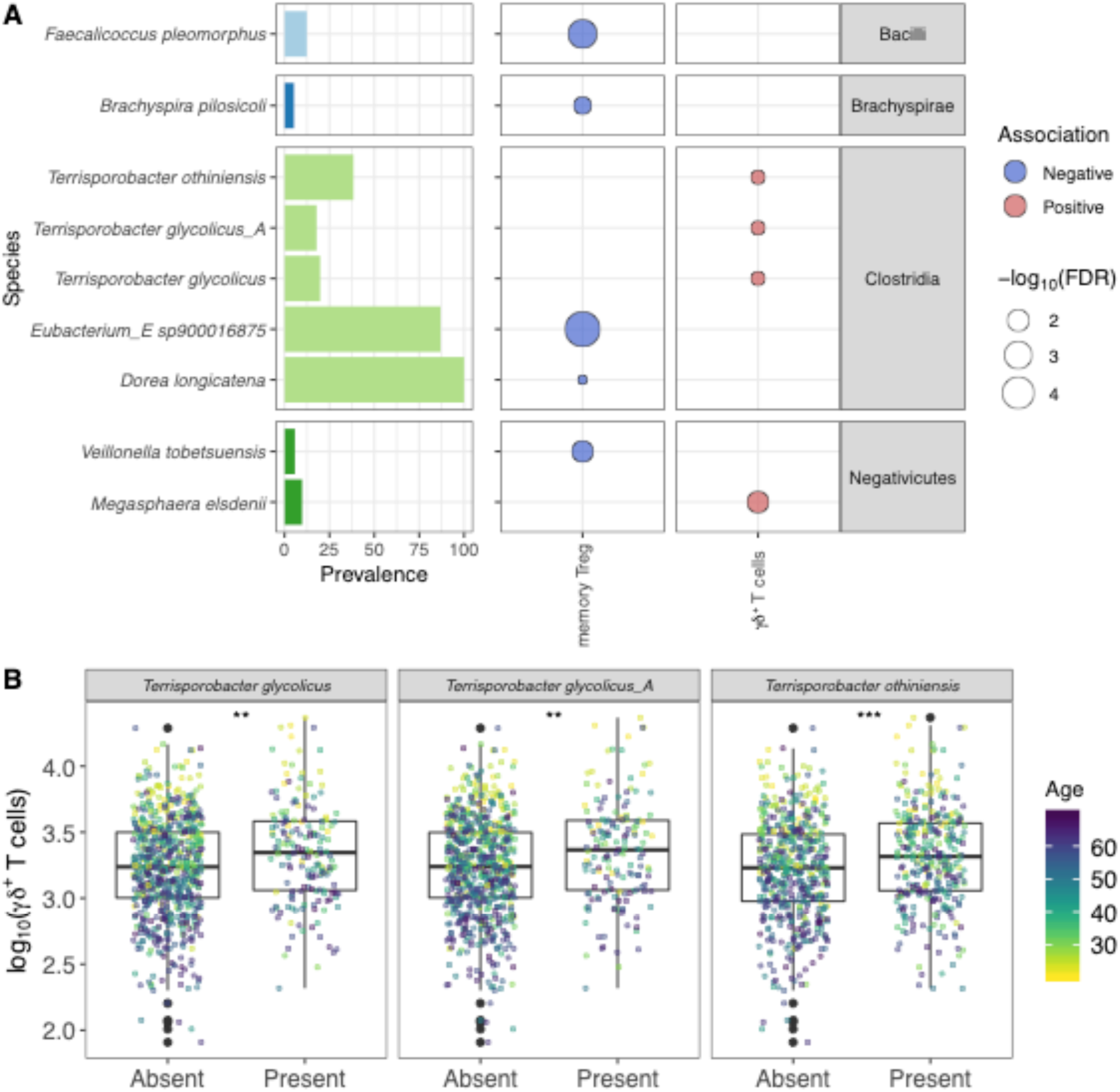
Bacteria species associated with circulating regulatory T (T_reg_) and γδ^+^ T cells, Related to Figure 5. (A) Right bars indicate the prevalence of the species in the MI donors and are grouped by bacterial class. In the center, a circle represents each significant immunophenotype x species interaction. Fill color indicates whether the interaction was positive/negative and size the -log10 FDR value. (B) Boxplots show the number of γδ^+^ T-cells in donors with or without the *Terrisporobacter* species. ** p < 0.01 by Wilcoxon-Rank Sum. See also Table S21.

**Figure S11.**
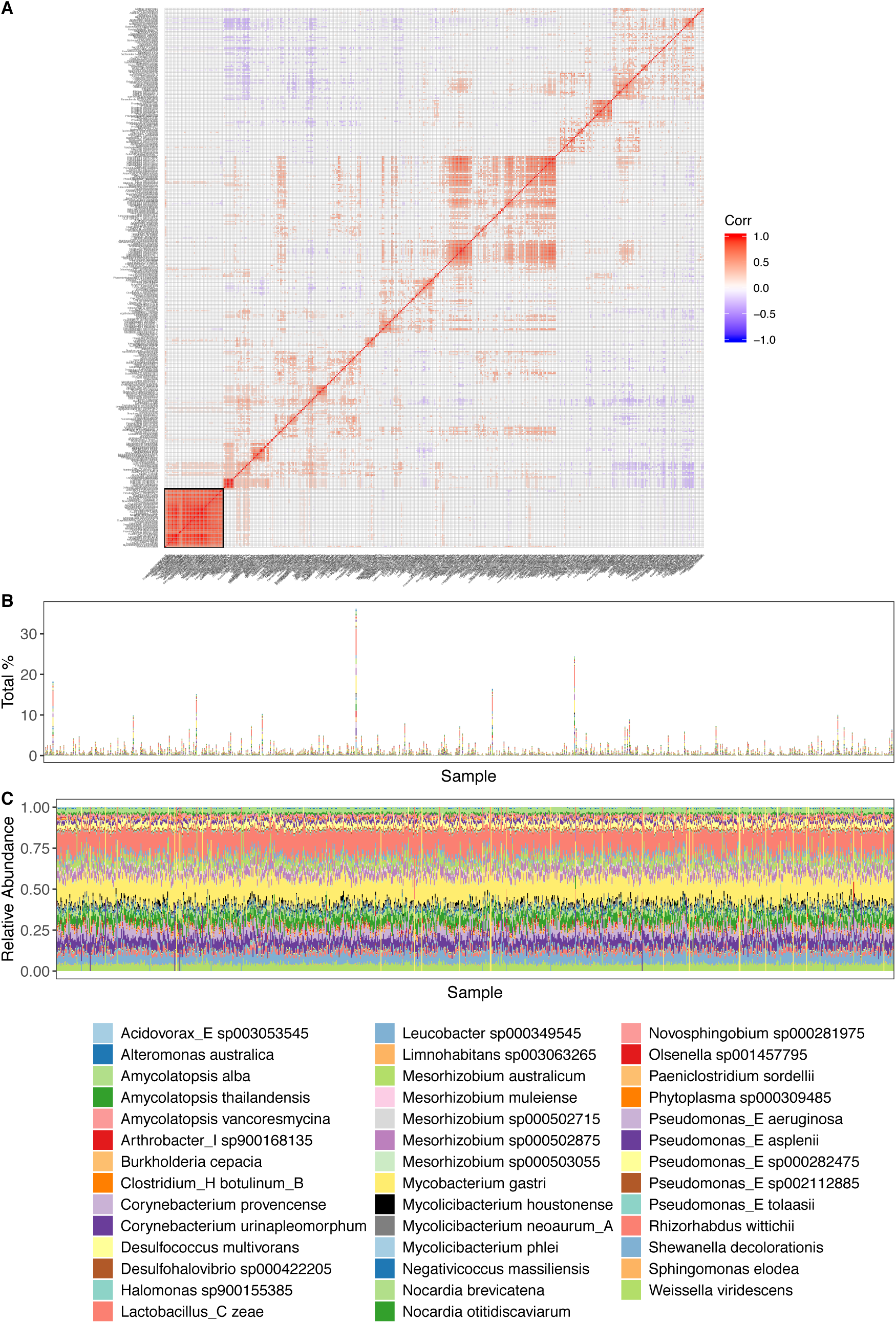
Putative reagent contaminants identified in the MI samples. (A) Correlation plot of 393 species in the MI donors with prevalence > 30%. Boxed are the 41 species identified as putative reagent contaminants. (B) Barplots show the portion of all microbial reads mapping to contaminant organisms. (C) Barplots show how the contaminant reads are distributed across species consistently across samples. See also Table S23.

**Figure S12.**
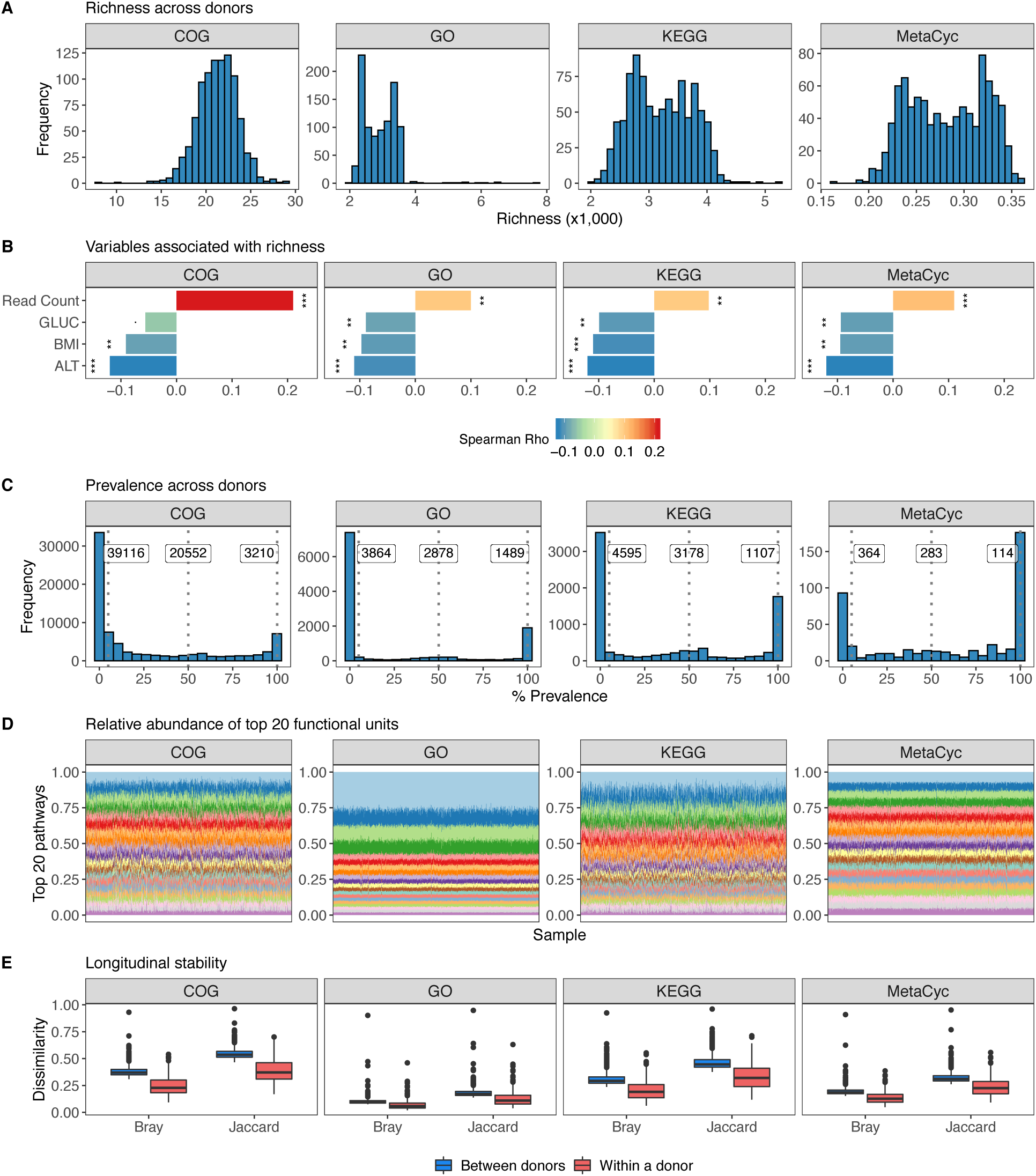
Microbial pathway profiles across donors. (A) Distribution of pathway/gene richness across donors and different functional databases. (B) Variables associated with functional richness based on Spearman. *** p-value < 0.001, ** p-value < 0.01. (C) Distribution of pathway prevalence across donors. The first line and value to the left indicates the number of pathways present in at least 5% of donors, the next 50%, and the final 100%. (D) Relative abundance plots of the top 20 functional units per database based on prevalence and mean abundance. (E) Boxplots of Bray-Curtis distances and binary Jaccard distances between donors and within a donor over time (n= 413, time between samples= 17 ± 3.3 days), 1 = samples are completely different, 0 = samples are identical. For all Wilcoxon-Rank Sum p-values were < 2.2e-16.

